# An energy-saving glasshouse film reduces seasonal, and cultivar dependent Capsicum yield due to light limited photosynthesis

**DOI:** 10.1101/2022.10.29.513818

**Authors:** Sachin G. Chavan, Xin He, Chelsea Maier, Yagiz Alagoz, Sidra Anwar, Zhong-Hua Chen, Oula Ghannoum, Christopher I. Cazzonelli, David T. Tissue

## Abstract

Glasshouse films can be used to reduce energy costs by limiting non-productive heat-generating radiation, but the impact on yield of greenhouse horticultural crops remains unknown. The effects of energy-saving film ULR-80 (referred to as Smart Glass; SG) designed to block long wavelength light that generates heat also reduced photosynthetically active radiation (PAR) consequently affecting crop morphology, photosynthesis, leaf pigments, and yield of two hydroponically grown capsicum (*Capsicum annuum* L.) cultivars (Red and Orange). The crops were grown in four high-tech glasshouse bays over two seasons of similar daily light integrals (DLI) during ascending (Autumn) and descending (Summer) photoperiods. The Red cultivar exhibited higher photosynthetic rates (light saturated - *A_sat_* and maximal - *A_max_*) and yield than the Orange cultivar in control but displayed stronger reductions in modelled photosynthetic rates at growth light and yield in SG without changes in photosynthetic capacity. Foliar pigment ratios of chlorophyll a/b and carotenoid: chlorophyll remained unaffected by the SG during both seasons indicating that chloroplast homeostasis was similar between SG and control. The seasonal differences in photosynthetic pigments and xanthophyll de-epoxidation state (DPS) revealed that cultivars were able to sense the SG-altered light environment during the ascending, but not descending photoperiod. The descending photoperiod correlated with a lower daily light level and a substantial yield reduction of 29 % and 13 % in Red and Orange cultivars, respectively. Thus, SG-induced higher reductions in yield during the descending photoperiod indicate that SG may be more beneficial for capsicum crops planted during Autumn with an ascending photoperiod.

**Highlights:**

- A potential energy saving SG film limited net photosynthesis of capsicum
- The SG film reduced yield of two capsicum cultivars that can be mitigated by planting during the low light growth season with a shorter photoperiod
- SG reduced genotype-dependent capsicum yield was associated with alterations in the level of foliar pigments required for photoprotection under adverse light conditions

## 1. Introduction

The global energy price is rising due to climate change and regional conflicts in recent years, and this will be an ongoing issue due to population growth and depletion of important non-renewable energy sources (Tollefson, 2022). The protected cropping industry requires research and innovation to contribute to sustainable food production (O’Sullivan et al., 2019), particularly to minimise energy use and operational costs in high tech greenhouses (Chavan et al., 2020; Shen et al., 2018). Energy saving cover materials may have a positive (Lemarié et al., 2019; Shen et al., 2021), minimal (He et al., 2021) or even negative (Chavan et al., 2020; Ntinas et al., 2019) impact on crop yield and quality, depending on the light transmission properties of the cover material, growth season, crop and cultivar (Loik et al., 2017; Timmermans et al., 2020). Studies investigating the impact of cover materials are not generally conducted at a commercial scale (Loik et al., 2017; Shen et al., 2021) nor do they comprehensively assess plant growth responses to altered light regimes caused by seasonal change or reduction in PAR (Aroca-Delgado et al., 2019). Recently, we have investigated the impact of light changes on crop growth, photosynthesis, yield, post-harvest fruit quality, and energy assessment using commercial practices of greenhouse vegetable production of eggplant and capsicum in a research facility with the precise climate control of a high-tech commercial glasshouse (Chavan et al., 2020; He et al., 2021, 2022; Lin et al., 2022; Zhao et al., 2021).

Our understanding of the role of light in plant growth and development is constantly evolving with ongoing research (Davis and Burns, 2016; Dou et al., 2017; Smith et al., 2017; Zhen and Bugbee, 2020). Ultraviolet (UV < 400 nm), PAR (400–700 nm), and near-infrared (NIR, 700–2500 nm) are the most relevant light wavelengths for greenhouses (Timmermans et al., 2020). Photon ratios affecting plant photobiology vary on a daily and seasonal basis (Kotilainen et al., 2020), which are further altered by seasonal variation in light transmission under the cover films. Light transmission through cover materials may change in intensity or spectral quality due to a partial or complete blockage of selective wavelengths (Chavan et al., 2020; Loik et al., 2017), or may increase in specific light wavelengths due to spectral shifting properties of the film (Shen et al., 2021). Light-altering cover materials can impact all aspects of development, including germination, vegetative growth, flowering and fruit ripening (Jones, 2018), partly dependent upon the season influence (Cerny et al., 2003). A study investigating influence of photo-selective films and growing season found that the far-red light absorbing films delayed anthesis and overall development of the floral meristem was slower than control during weakly inducting photoperiod (Cerny et al., 2003). The seasonal influence of light altering cover films on plant productivity in glasshouses dependent upon ascending and descending photoperiods during winter and summer periods, respectively, remains under studied in the protected cropping environment.

The time of planting plays an important role in crop growth, development and yield particularly due to changes in photoperiod (Dorais et al., 1996). Longer photoperiods with the same DLI can improve plant growth via increased plant dry mass, height, leaf area, and chlorophyll content (Elkins and Iersel, 2020). Furthermore, photoperiod positively or negatively affects plant performance depending on the plant developmental stage. For example, longer photoperiod during the reproductive stage may increase yield due to higher carbon translocation rates (Dorais et al., 1996) and shorter photoperiods may favour seedling growth (Yang et al., 2017). Source (supply) and sink (demand) regulation maintains balance between vegetative and reproductive growth by aborting reproductive organs (Wubs et al., 2009) and affects fruit set, leading to periods of high yield alternating with periods of low yield for indeterminate plants, such as capsicum (Ma et al., 2011). Thus, low light limits photosynthesis and yield (Chavan et al., 2020; Hao and Papadopoulos, 1999) mainly through a lower supply of carbon from source leaves to sinks, such as fruits (Aloni et al., 1996; Li et al., 2015; Turner and Wien, 1994).

The chloroplast is both a sensor and integrator of environmental information, playing a crucial role in the process of photo-acclimation in leaves (Dhami and Cazzonelli, 2020). Plant leaves utilise chlorophyll pigments to capture light energy and drive photosynthesis in the photosystems and incorporate carotenoids as accessory pigments to dissipate excess light via chlorophyll *a* fluorescence, as heat (nonphotochemical quenching), or spent on the transition of Chl to a triplet state that generates singlet oxygen and may cause destruction of the photosynthetic apparatus (Baker, 2008; Demmig-Adams et al., 2014). Non-photochemical dissipation of excess light using the xanthophyll cycle pigments involves de-epoxidation of light-harvesting violaxanthin to the energy quenching antheraxanthin and zeaxanthin photoprotective pigments (Demmig-Adams et al., 2014). When plants are exposed to high light, extreme heat waves, or an altered light environment, the foliar xanthophyll DPS changes to maintain chloroplast homeostasis and photosynthesis capacity (Chavan et al., 2020; Dhami et al., 2020). Seasonal changes in photoperiods and shading were shown to alter foliar pigment composition and the DPS (Garcia-Plazaola et al., 1997). Nonetheless, it remains unclear how a SG film and seasons will impact the DPS of different cultivars within a protected cropping environment.

SG film was designed to block heat-generating long wavelength light, while allowing transmission of light useful for photosynthesis and plant growth. It was previously tested in eggplants grown for six months, primarily in high light conditions of spring and summer (Chavan et al., 2020). SG decreased energy use and increased nutrient- and water-use efficiency, but also reduced eggplant fruit yield due to light-limited photosynthesis and consequent source-sink regulated fruit abortion. Here, we assess the seasonal impact of SG on capsicum crop yield during periods of ascending photoperiod (Autumn Experiment; AE) and descending photoperiod (Summer Experiment; SE), but with similar DLI over the entire growth period. We compare two cultivars (Red and Orange) of capsicum in two experimental trials (38 weeks per trial) that utilise standard growth management practices. The impacts of SG on capsicum growth and production address the following hypotheses: SG will; 1) reduce light and hence photosynthetic rates and capsicum yield; 2) reduce the yield of capsicum to a greater degree during the descending photoperiod than the ascending photoperiod, as the DLI is reduced during crop establishment; and 3) alter foliar pigment levels and/or ratios to maintain chloroplast homeostasis and light energy harvesting via reduction in photoprotective pigments (Xanthophyll DPS) particularly during high light seasons (Figure 1).

**Figure 1.**
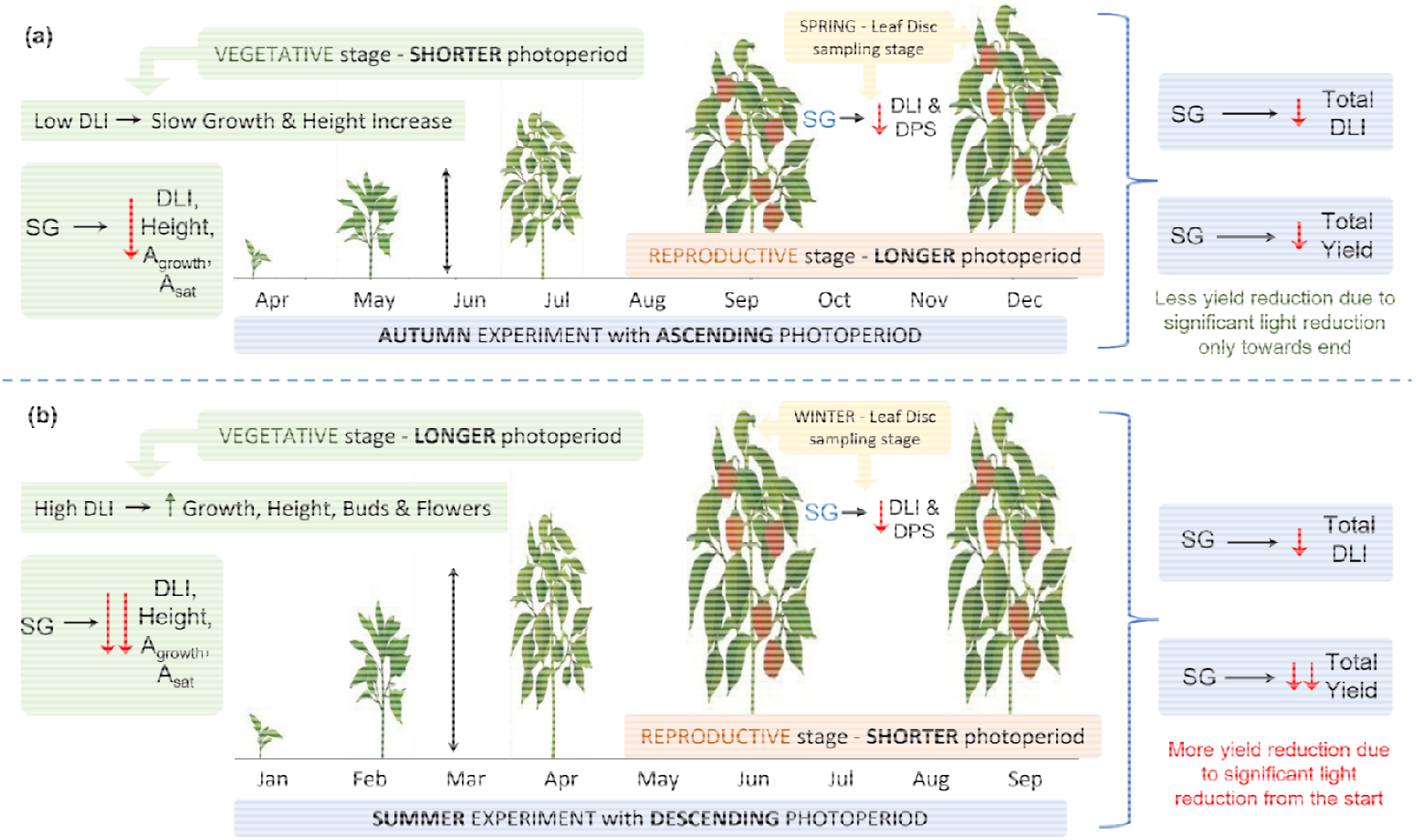
Hypotheses for effects of Smart Glass (SG) during Autumn Experiment (AE) with Ascending photoperiod (a) and Summer Experiment (SE) with Descending photoperiod (b) o Capsicum cultivars Red and Orange. High light during crop establishment and vegetative growth in SE will promote plant growth and development relative to AE. Based on eggplant trial is predicted to reduce daily light integral (DLI), light saturated photosynthesis (A_sat_) and net photosynthesis at growth light (A_growth_) more during SE with high light levels at the st experiment than AE. We predict that despite similar reductions in total growth season DLI under SG in both experiments, total yield be more severely reduced in SE due to higher reduct DLI during crop establishment. SG will alter foliar pigments by reducing xanthophyll pigments involved photoprotection. Up and down arrows indicate increase decrease in the mea traits.

## 2. Materials and Methods

### 2.1. Plant material and experimental design

The capsicum trial was conducted in a high-tech greenhouse facility with east-west orientation located on the Hawkesbury Campus of Western Sydney University, Richmond, NSW, Australia. The facility included four experimental bays (105 m^2^ each) fitted with HD1AR diffuse glass on the roof (70% haze) and on the sidewall (5% haze). Two bays were coated with SG film (Solar Gard, Saint-Gobain Performance Plastics, Sydney, Australia) on both roof and side walls and the other two bays were used as “Control’ as described previously (Chavan et al. 2020; Zhao et al. 2021). To investigate the effect of SG and time of planting on capsicum, a 2×2 factorial design was adopted with two cultivars, Orange (O06614 in the first trial and Kathia in the second trial, respectively) and Red (Gina, Syngenta, USA). The two experimental trials were conducted from 19 April 2019 to 19 December 2019 (AE with ascending photoperiod) and from 20 January 2020 to 20 September 2020 (SE with descending photoperiod).

The capsicum cultivars, Red and Orange, were sown into Rockwool cubes (Grodan, The Netherlands) and plants were transplanted 42 days after sowing on the19^th^ April 2019 (AE) in the first trial and 20^th^ January 2020 (SE) in the second trial. Each research bay (105 m^2^) included six gutters that were 10.8 m (length) × 25 cm (width, AIS Greenworks, Castle Hill, NSW, Australia), and contained 10 Rockwool slabs (90×15×10 cm, Grodan, The Netherlands) as a substrate for plants, per gutter. Plants were grown under non-limiting water and nutrient (EC: 2.5~3.0 dS m^−1^, pH: 5.0-5.5) conditions, at [CO_2_] (489.6 and 476.6 μl l^−1^ daytime average), temperature (25.3/19.3 and 25.2/19.3 °C day/night average), relative humidity (RH; 74.2/72.9 and 74.2/77.5 %, day/night average) and natural light in AE and SE with ascending and descending photoperiod, respectively, according to the commercial practices of vertical hydroponic production of fruits and vegetables in greenhouses. Day and night-time averages were calculated from 10.00 h to 16.00 h and from 20.00 h to 06.00 h, respectively. Individual capsicum cultivars were alternatively transplanted in each gutter so that two of the middle four gutters contained the Red cultivar and the other two middle gutters contained the Orange cultivar. The two side gutters were planted with the Red cultivar, which were not used for measurements to account for the edge effect, and measurements were performed on plants in the middle four gutters. Similar crop management practices of cutting and pruning were used for cultivation in both experiments. Each stem was considered as an individual replicate, and all measurements were performed on a per stem basis. Both experiments lasted for 38 weeks after transplanting.

### 2.2. Growth environment measurements

The environmental parameters were monitored and controlled by Priva software and hardware (Priva, The Netherlands). Priva-controlled growth environment parameters included atmospheric CO_2_, air temperature and RH at canopy level, and PAR at roof level. An additional five PAR sensors (190R-SMV-50 Quantum Sensor, LI-COR) were distributed spatially across each bay to measure light at canopy level during both experimental trials. Net radiometer (SN-500, Apogee Instruments) and diffuse light sensors (BF5 sunshine sensor, Delta T Devices) were used to measure fluxes of net shortwave-longwave radiation and light diffusion, respectively. The Priva system and sensor operation is fully described in (Chavan et al., 2020; Lin et al., 2022).

### 2.3. Growth, morphology and productivity measurements

Plant growth was measured on five plants (each stem refers to individual plant hereafter) per gutter in each room (n=20 plants per cultivar in the Control or SG room) in both experimental trials. The plant height was measured every fortnight. The number of buds, flowers and developing fruits in the branch were recorded once a week from five plants per gutter in each room (n=20 per cultivar in Control or SG), and the average bud and number of flowers were calculated per stem (n stem^−1^). Mature fruits, selected based on the colour (red and orange) of individual fruits, were harvested, and the individual fruit weight and the number of fruits per stem were recorded every week. The fruits were graded as marketable (≥ 100 g, including the extra-large fruit ≥ 250 g) and unmarketable fruits, which included small (< 100g, edible) fruits and fruits with rotting, cracking, lobing and other deformities.

### 2.4. Leaf gas exchange measurements

Instantaneous steady-state leaf gas exchange measurements were performed using a portable, open-mode gas exchange system (LI-6400XT, LI-COR, Lincoln, USA) during ascending and descending photoperiod in September 2019 and March 2020, respectively. Photosynthetic traits were measured at saturating light of 1500 μmol m^−2^ s^−1^ PAR, 500 μl L^−1^ CO_2_ concentration and 25°C leaf temperature: *A*_sat_, stomatal conductance (*g*_s_), the ratio of intercellular to ambient CO_2_ (*C*_i_/*C*_a_), transpiration rate (E, mmol m^−2^ s^−1^), dark respiration (R_d_), and photosynthetic water use efficiency (PWUE) calculated as A_sat_/*g*_s_. Measurements were taken between 10 am and 2 pm on the first fully expanded leaf of the main stems. The response of *A_sat_* to light (Q) (A-Q curve) was measured at 25°C leaf temperature at nine light levels (0, 25, 50, 100, 250, 500, 1000, 1500 and 2000 μmol m^−2^ s^−1^) during ascending photoperiod and descending photoperiod (n = 12 per cultivar in Control or SG room) in September 2019 and March 2020, respectively. Dark respiration (R_d_) was measured by switching the light off for 15-20 min during the dark adaptation period. Maximum light-saturated CO_2_ assimilation rate (*A_max_*), the maximum quantum yield of PSII (*ϕ*_max_) and curvature factor of the light response curve (θ) were measured to assess the effects of SG. *ϕ*_max_ was also determined using the initial slope of the light response curves. The light response curve means were fitted using the following equation (Ögren and Evans, 1993; Xu et al., 2019).

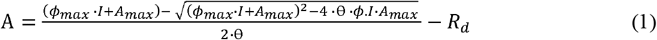

where, *I* = absorbed irradiance, we assumed absorptance = 0.85; *A* = CO_2_ assimilation rate at given light; R_d_ = dark respiration; Φ_max_ = maximum quantum yield of PSII; *A*_max_ = maximum light-saturated CO_2_ assimilation rate; and θ = curvature factor of the light response curve.

To determine A at growth light in Control and SG, average canopy PAR (measured using five sensors at canopy level from 9 am to 3 pm) was used to model CO_2_ assimilation rates (*A_model_*) using light response curves according to equation (1).

### 2.5. High-performance liquid chromatography analysis of leaf pigments

Carotenoids and chlorophyll analysis was performed on one leaf per plant (n=10) using high-performance liquid chromatography (HPLC) (Agilent 1260 Infinity, Agilent, Santa Clara, CA, USA). Fully expanded mature leaves were punched from the middle position to collect leaf discs using a size 10 (2.54 cm^2^ leaf area) cork-borer. Samples were snap-frozen using liquid nitrogen and stored at −80 °C. Leaf mass per unit area (LMA) and carotenoids and pigments were quantified using fresh leaf weight. Frozen Samples were ground to a fine powder with TissueLyser (Qiagen, Germany) and pigments extracted as previously described (Alagoz et al., 2020). Carotenoids and chlorophylls were identified using their retention time and light emission absorbance spectra, and absolute quantification was performed as described earlier (Anwar et al., 2022). The xanthophyll cycle de-epoxidation (violaxanthin;V, antheraxanthin;A. zeaxanthin;Z) was determined as the ratio of = (A+Z)/(A+Z+V), which we herein refer to as the DPS.

### 2.6. Statistical analysis

Statistical data analyses were performed using R statistical programming software (https://www.r-project.org/, 4.2.1, R Core Team, 2022). Linear regression was used to examine the bivariant relationship among parameters over SG and Control. *P* ≤ 0.05 was considered as statistical significance. Levene’s test from the *car* package was used to test the homogeneity of variance. Welch’s t-test for unequal variances was used for parameters showing unequal variance. Shapiro-Wilk test for normality was used to test the normal distribution. Parameters with non-normal distribution were then analysed using a non-parametric equivalent of one-way anova (Kruskal-Wallis Test). Other packages included *lubridate* (for effective use of dates in plots), *sciplot* (for plotting), *tidyverse and dplyr* (for data manipulation), *HIEv* (for data download from local sever), and *doBy* (for calculating means and standard errors). The effects impact of SG, experiments, cultivars and their interactions on plant growth, yield and quality parameters were analysed using one-way and two-way analysis of variance (ANOVA) functions in *car* package. Linear modelling function in R was used for correlation analysis. The significance levels were ns, *, **, and *** indicated *P* > 0.05, *P* ≤ 0.05, *P* ≤ 0.01, and *P* ≤ 0.001, respectively.

## 3. Results

### 3.1. SG significantly decreases PAR except in Winter low light conditions with shorter photoperiod

The SG film altered the light quality, reducing UV (221~400 nm, −69%, *P* ≤ 0.001), red (600~700 nm, −26%, *P* ≤ 0.01) and far-red (700~800 nm, −51%, *P* ≤ 0.05) radiation (Table S1). SG significantly reduced the DLI during the higher light seasons in both ascending and descending photoperiod experiments: AE and SE (Figure 2). Although both experiments covered all four seasons, plants were established during different light conditions. The AE experiment, with ascending photoperiod, started in Autumn under moderate DLI (~10 mol m^−2^ d^−1^) and a short photoperiod (Figure 2-C), followed by low DLI during Winter (~8.5 mol m^−2^ d^−1^ in Control, Table 1-C), without significant light differences between SG and Control. SG reduced DLI in Spring (~20 mol m^−2^ d^−1^ in Control) by 21% (*P* ≥ 0.001) during the second half of the AE experiment. The SE experiment, with descending photoperiod, started in Summer under high light conditions (DLI ~15 mol m^−2^ d^−1^ in Control) and a long photoperiod (Figure 2-C), with SG reducing DLI by 21% (*P* ≥ 0.001) during the first half of the SE experiment. SG had no significant effect on DLI during the subsequent Winter (DLI ~8.1 mol m^−2^ d^−1^ in Control, Table S2), but reduced light during the Spring (DLI ~ 20 mol m^−2^ d^−1^ in Control), at the end of the SE experiment (Figure 2). Overall, SG reduced average growth season DLI slightly more during descending (−22%, *P* ≤ 0.001) than ascending photoperiod (−18%, *P* ≤ 0.001). However, SG reduced DLI to a greater extent during high light (−24%, *P* ≤ 0.001) periods of Spring through Summer (August to April) relative to low light periods (−6%, *P* ≤ 0.07) of peak Winter months (May to July) during both ascending and descending photoperiods (Figure 2, Table S2). Thus, compared to AE with ascending photoperiod, SE with descending photoperiod received relatively less light which was further reduced under SG more in SE, particularly during high light conditions.

**Figure 2.**
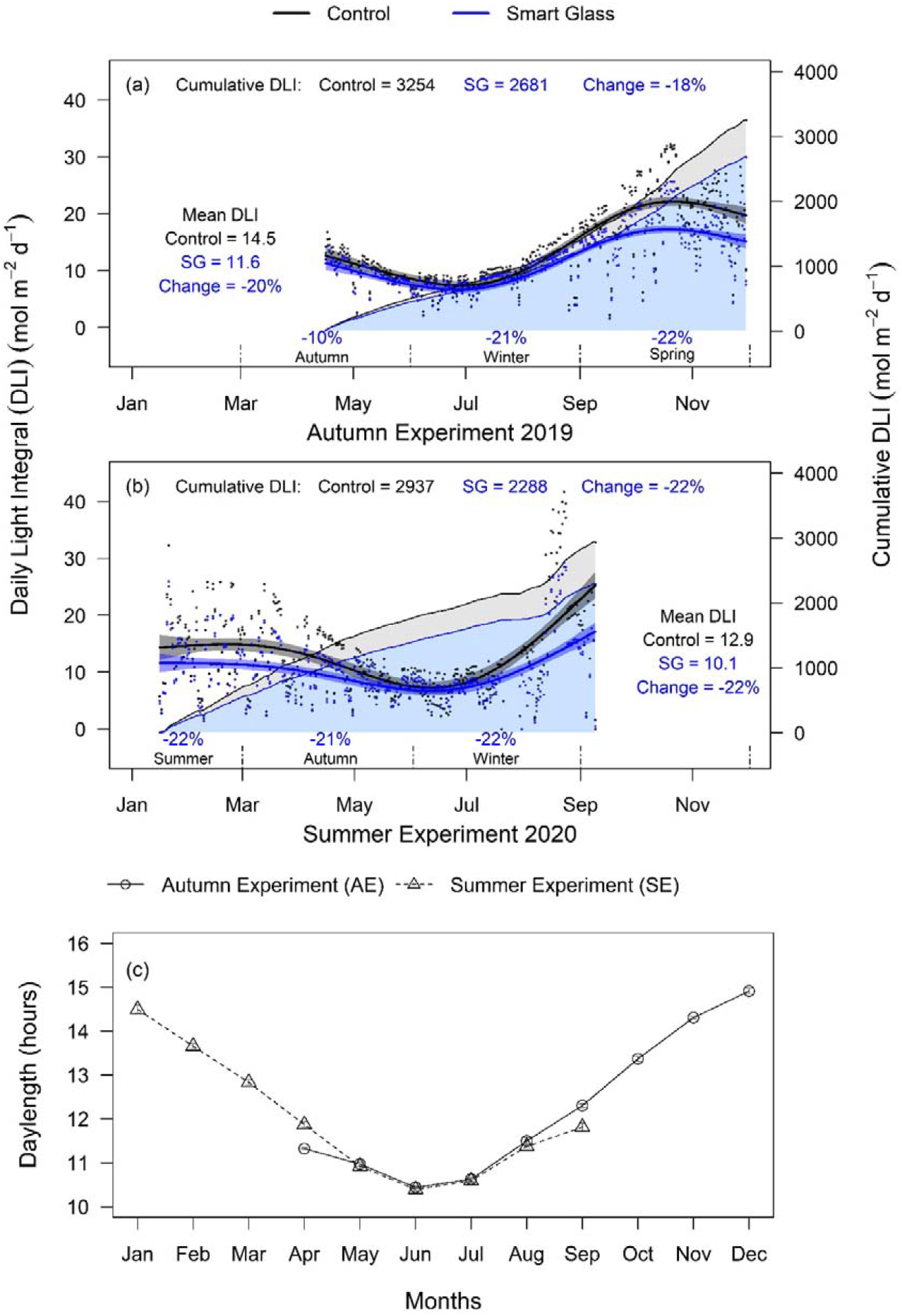
Smart Glass (SG) impact on light transmittance in two experimental trials. The smooth plot of daily light integral (DLI, total daily PAR) and cumulative DLI over time are presented for (a) Autumn Experiment (AE, 19 April to 19 Dec 2019) with ascending photoperiod and (b) Summer Experiment (SE, 20 Jan to 20 Sept 2020) with descending photoperiod. Points indicate average DLI measured using five PAR sensors at canopy level. Control (black) and SG (blue) treatments are depicted with 95 % confidence intervals. The % difference in mean experimental DLI (SG/Control) is indicated for each season. Line plot in panel (c) depicts the monthly means of photoperiod with standard error during AE (circles) and SE (triangles).

**Table 1.**
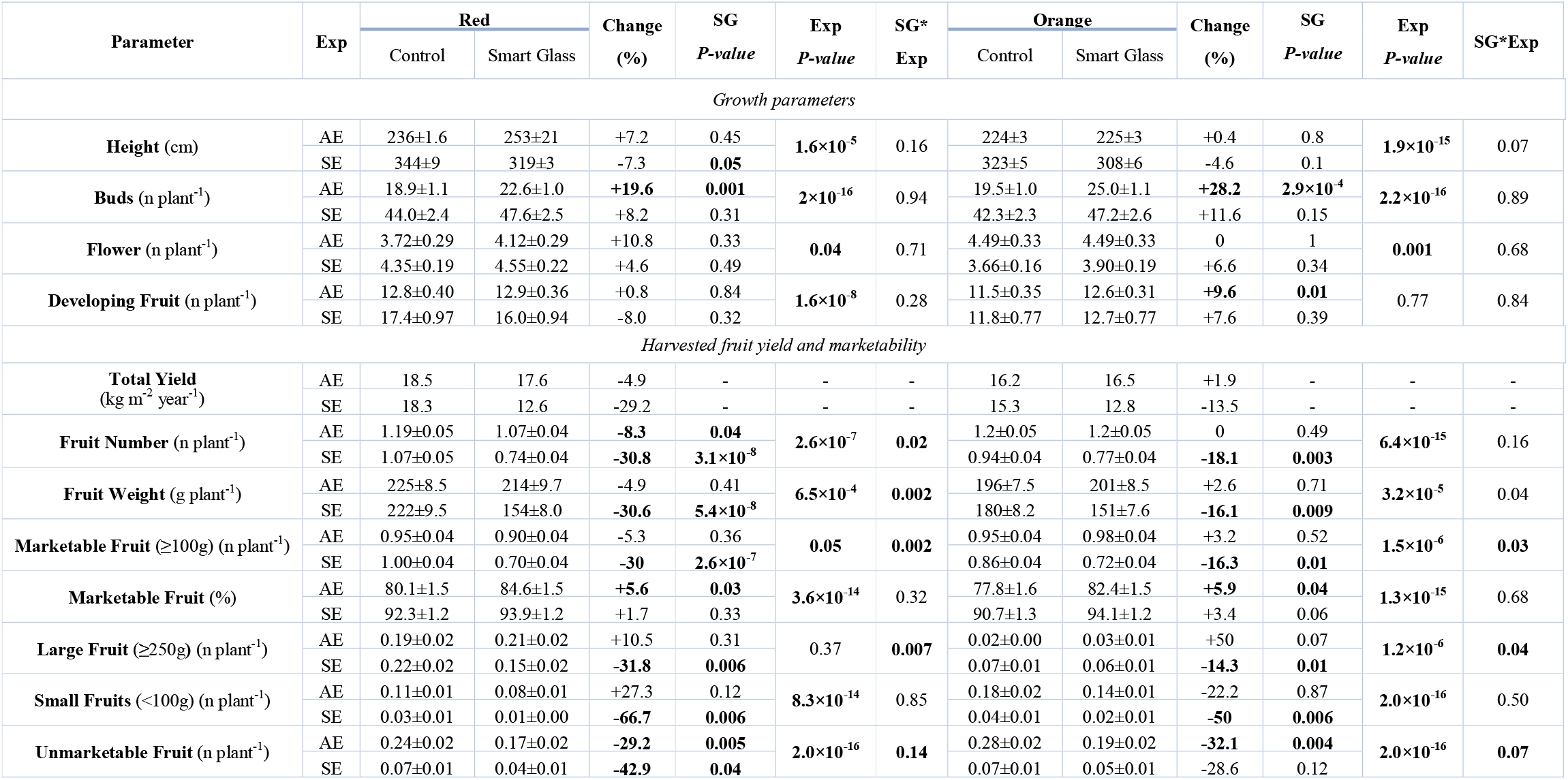
Summary of statistical analyses for the SG and experiment effect on growth and productivity parameters. The values are mean ± standard error of mean (n = 20-40). Change represents SG/Control. *P* values are given according to the one-way analysis of variance (ANOVA) (SG), two-way ANOVA (SG*Exp) or Welch’s F-test for equal and unequal variance respectively and *P* ≤ 0.05 are significant. For non-normal distribution Kruskal-Wallis non-parametric test was used. AE – Autumn Experiment with ascending photoperiod, SE – Sumer Experiment with descending photoperiod.

| Parameter | Exp | Red |  | Change<br>(%) | SG<br>*P-value* | Exp<br>*P-value* | SG*<br>Exp | Orange |  | Change<br>(%) | SG<br>*P-value* | Exp<br>*P-value* | SG*Exp |
| --- | --- | --- | --- | --- | --- | --- | --- | --- | --- | --- | --- | --- | --- |
|  |  | Control | Smart Glass |  |  |  |  | Control | Smart Glass |  |  |  |  |
| Growth parameters |  |  |  |  |  |  |  |  |  |  |  |  |  |
| Height (cm) | AE | 236±1.6 | 253±21 | +7.2 | 0.45 | 1.6×10<sup>-5</sup> | 0.16 | 224±3 | 225±3 | +0.4 | 0.8 | 1.9×10<sup>-15</sup> | 0.07 |
|  | SE | 344±9 | 319±3 | -7.3 | 0.05 |  |  | 323±5 | 308±6 | -4.6 | 0.1 |  |  |
| Buds (n plant<sup>-1</sup>) | AE | 18.9±1.1 | 22.6±1.0 | +19.6 | 0.001 | 2×10<sup>-16</sup> | 0.94 | 19.5±1.0 | 25.0±1.1 | +28.2 | 2.9×10<sup>-4</sup> | 2.2×10<sup>-16</sup> | 0.89 |
|  | SE | 44.0±2.4 | 47.6±2.5 | +8.2 | 0.31 |  |  | 42.3±2.3 | 47.2±2.6 | +11.6 | 0.15 |  |  |
| Flower (n plant<sup>-1</sup>) | AE | 3.72±0.29 | 4.12±0.29 | +10.8 | 0.33 | 0.04 | 0.71 | 4.49±0.33 | 4.49±0.33 | 0 | 1 | 0.001 | 0.68 |
|  | SE | 4.35±0.19 | 4.55±0.22 | +4.6 | 0.49 |  |  | 3.66±0.16 | 3.90±0.19 | +6.6 | 0.34 |  |  |
| Developing Fruit (n plant<sup>-1</sup>) | AE | 12.8±0.40 | 12.9±0.36 | +0.8 | 0.84 | 1.6×10<sup>-8</sup> | 0.28 | 11.5±0.35 | 12.6±0.31 | +9.6 | 0.01 | 0.77 | 0.84 |
|  | SE | 17.4±0.97 | 16.0±0.94 | -8.0 | 0.32 |  |  | 11.8±0.77 | 12.7±0.77 | +7.6 | 0.39 |  |  |
| Harvested fruit yield and marketability |  |  |  |  |  |  |  |  |  |  |  |  |  |
| Total Yield<br>(kg m<sup>-2</sup> year<sup>-1</sup>) | AE | 18.5 | 17.6 | -4.9 | - | - | - | 16.2 | 16.5 | +1.9 | - | - | - |
|  | SE | 18.3 | 12.6 | -29.2 | - | - | - | 15.3 | 12.8 | -13.5 | - | - | - |
| Fruit Number (n plant<sup>-1</sup>) | AE | 1.19±0.05 | 1.07±0.04 | -8.3 | 0.04 | 2.6×10<sup>-7</sup> | 0.02 | 1.2±0.05 | 1.2±0.05 | 0 | 0.49 | 6.4×10<sup>-15</sup> | 0.16 |
|  | SE | 1.07±0.05 | 0.74±0.04 | -30.8 | 3.1×10<sup>-8</sup> |  |  | 0.94±0.04 | 0.77±0.04 | -18.1 | 0.003 |  |  |
| Fruit Weight (g plant<sup>-1</sup>) | AE | 225±8.5 | 214±9.7 | -4.9 | 0.41 | 6.5×10<sup>-4</sup> | 0.002 | 196±7.5 | 201±8.5 | +2.6 | 0.71 | 3.2×10<sup>-5</sup> | 0.04 |
|  | SE | 222±9.5 | 154±8.0 | -30.6 | 5.4×10<sup>-8</sup> |  |  | 180±8.2 | 151±7.6 | -16.1 | 0.009 |  |  |
| Marketable Fruit (≥100g) (n plant<sup>-1</sup>) | AE | 0.95±0.04 | 0.90±0.04 | -5.3 | 0.36 | 0.05 | 0.002 | 0.95±0.04 | 0.98±0.04 | +3.2 | 0.52 | 1.5×10<sup>-6</sup> | 0.03 |
|  | SE | 1.00±0.04 | 0.70±0.04 | -30 | 2.6×10<sup>-7</sup> |  |  | 0.86±0.04 | 0.72±0.04 | -16.3 | 0.01 |  |  |
| Marketable Fruit (%) | AE | 80.1±1.5 | 84.6±1.5 | +5.6 | 0.03 | 3.6×10<sup>-14</sup> | 0.32 | 77.8±1.6 | 82.4±1.5 | +5.9 | 0.04 | 1.3×10<sup>-15</sup> | 0.68 |
|  | SE | 92.3±1.2 | 93.9±1.2 | +1.7 | 0.33 |  |  | 90.7±1.3 | 94.1±1.2 | +3.4 | 0.06 |  |  |
| Large Fruit (≥250g) (n plant<sup>-1</sup>) | AE | 0.19±0.02 | 0.21±0.02 | +10.5 | 0.31 | 0.37 | 0.007 | 0.02±0.00 | 0.03±0.01 | +50 | 0.07 | 1.2×10<sup>-6</sup> | 0.04 |
|  | SE | 0.22±0.02 | 0.15±0.02 | -31.8 | 0.006 |  |  | 0.07±0.01 | 0.06±0.01 | -14.3 | 0.01 |  |  |
| Small Fruits (<100g) (n plant<sup>-1</sup>) | AE | 0.11±0.01 | 0.08±0.01 | +27.3 | 0.12 | 8.3×10<sup>-14</sup> | 0.85 | 0.18±0.02 | 0.14±0.01 | -22.2 | 0.87 | 2.0×10<sup>-16</sup> | 0.50 |
|  | SE | 0.03±0.01 | 0.01±0.00 | -66.7 | 0.006 |  |  | 0.04±0.01 | 0.02±0.01 | -50 | 0.006 |  |  |
| Unmarketable Fruit (n plant<sup>-1</sup>) | AE | 0.24±0.02 | 0.17±0.02 | -29.2 | 0.005 | 2.0×10<sup>-16</sup> | 0.14 | 0.28±0.02 | 0.19±0.02 | -32.1 | 0.004 | 2.0×10<sup>-16</sup> | 0.07 |
|  | SE | 0.07±0.01 | 0.04±0.01 | -42.9 | 0.04 |  |  | 0.07±0.01 | 0.05±0.01 | -28.6 | 0.12 |  |  |

### 3.2. SG reduces crop height during the descending photoperiod associating with faster growth, increased bud and flower numbers

Higher DLI and longer daylength in the SE, with descending photoperiod, promoted faster growth during early seedling growth when compared to the AE, with ascending photoperiod. At 32 days after transplanting (DAT), both descending and ascending photoperiod capsicum plants had similar height. However, plants grown during the descending photoperiod were 44% (at 70 DAT) and 22% (at 185 DAT) taller than ascending photoperiod counterparts in both Control and SG treatments (Table 1). During the early growth period, SG reduced (−5%, *P* ≤ 0.05) plant height relative to the control, but the final plant height was similar under both treatments and experiments (Figure 3, a-b, Table 1).

**Figure 3.**
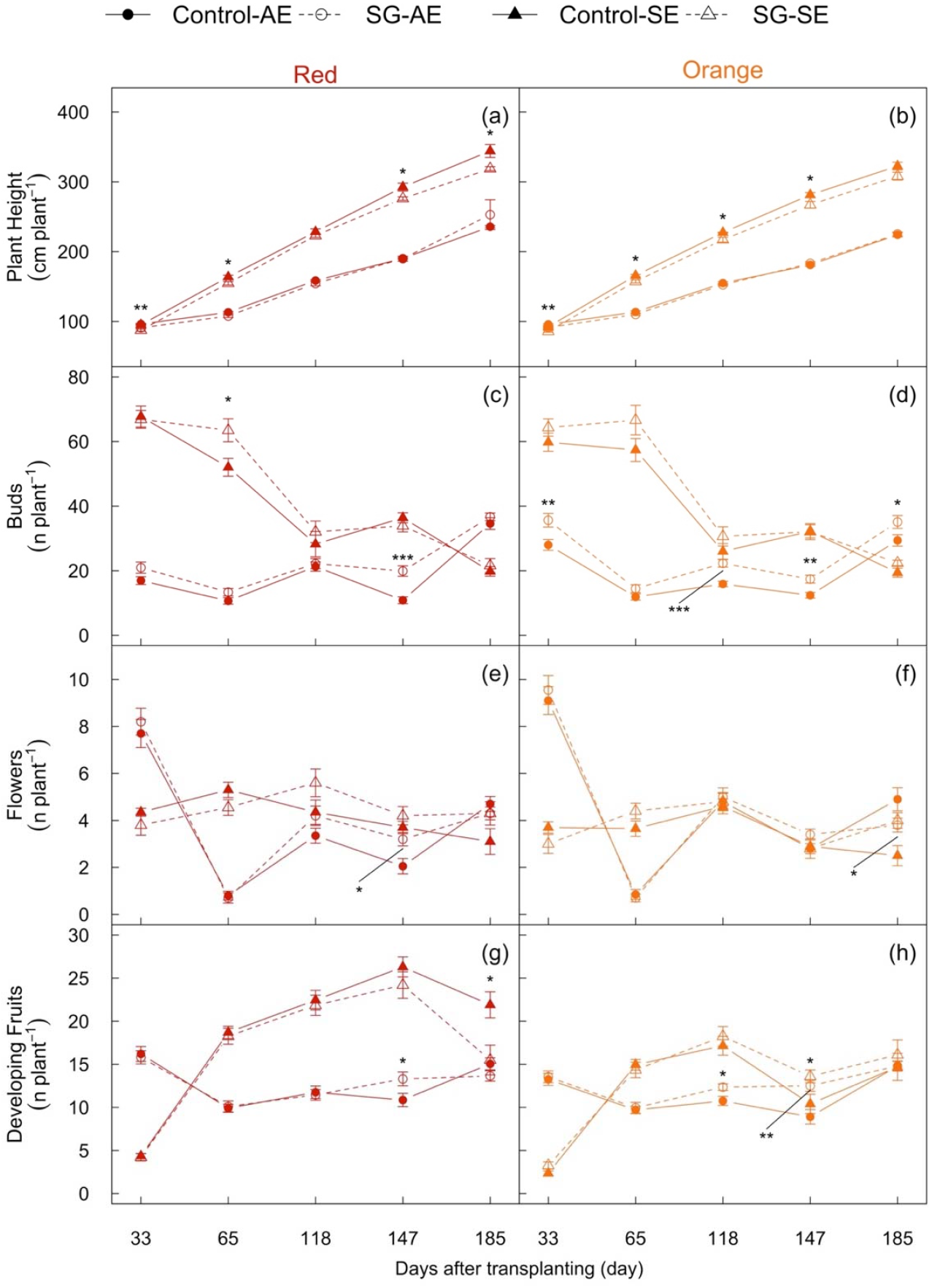
Smart Glass (SG) impact on morphological parameters in Red and Orange capsicum cultivars during Autumn (AE) and Summer (SE) experiment. Panels a and b depict cumulative stem height, c and d depict number of buds, e and f depict number of flowers and g and h depict number of developing fruits over time. Control and SG treatments are depicted in solid and dashed lines, respectively. Circles and triangles represent Autumn and Summer experiment, respectively. Error bars indicate standard error (n=20) of mean. The dashed and solid arrows indicate the time of first harvest in Autumn and Summer experiment, respectively. Statistical significance levels (*t*-test) for SG effect are shown: *, *P* ≤ 0.05; **, *P* ≤ 0.01: ***, *P* ≤ 0.001.

Bud number was higher (+86%, *P* ≤ 0.001) during descending relative to ascending photoperiod, and SG increased the bud number for both the Red (+20% *P* ≤ 0.001) and Orange (+28% *P* ≤ 0.001) cultivars only during the ascending photoperiod (Figure 3, c-d, Table 1). Both cultivars maintained an average of four flowers per stem (Figure 3, e-f, Table 1). In general, SG did not affect mean bud number except for a few instances such as at 147 DAT. SG increased the mean flower number in the Red cultivar by 56% during ascending photoperiod (Figure 3, Table 1). The number of developing fruits was generally higher in descending relative to ascending photoperiod for both cultivars, but significantly higher (+27%, *P* ≤ 0.001) in the Red cultivar. SG did not affect the number of developing fruits, but an increase was observed (+9.6 %, *P* ≤ 0.01) for the Orange cultivar during the ascending photoperiod (Figure 3, g-h, Table 1).

Taken together, both cultivars grew slower (evident from lower height and reproductive organs including buds, flowers and developing fruits) during AE with ascending photoperiod due to the lower light and daylength during early crop establishment. SG promoted plant growth and development by increasing reproductive organs in SE with descending photoperiod, but SG reduced height when plants were actively growing in SE due to high light availability.

### 3.3. SG reduces yield more during descending than ascending photoperiods yet increases the number of marketable fruits

Capsicum yield was determined using weekly harvested fruit number and weight; the first harvest occurred at 61 DAT and 79 DAT in ascending and descending photoperiod, respectively (Figure 3). SG reduced yield by lowering DLI, but the extent varied with cultivar, experiment, and season (Figure 4 and 5, Table 1). During the ascending photoperiod, SG decreased mean fruit number in the Red cultivar (−8 %*P* ≤ 0.04, Table 1), mainly at the end of growth season due to significant light differences in the second half of the growth season (Figure 4, a, c, e and Figure 5). SG significantly reduced light intensity during the descending photoperiod at the start of the growing season (Figure 4, b), which led to a reduction in fruit number and weight in the Red (−31%, *P* ≤ 0.001 and −30%, *P* ≤ 0.001, respectively) and Orange (−18%, *P* = 0.003 and −16%, *P* = 0.009, respectively) cultivars (Figures 4-b, d, f and 5, Table 1).

**Figure 4.**
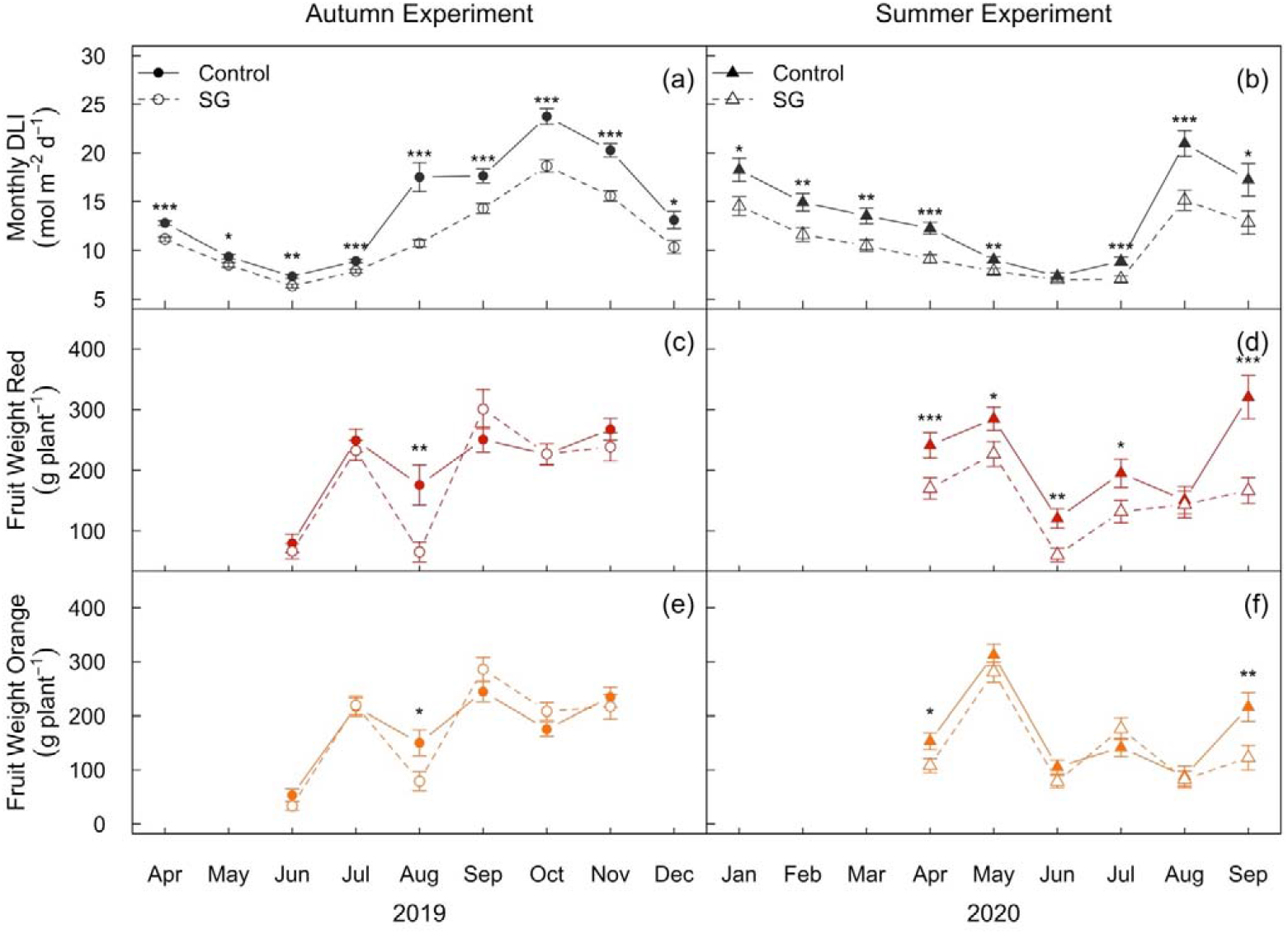
Impact of Smart Glass (SG) on monthly averages of daily light integrals (DLI) and harvested fruit weight across the growth season. Panel a and b depict the monthly means for canopy level DLI during Autumn Experiment (AE, Ascending Photoperiod) and Summer Experiment (SE, Descending Photoperiod), respectively. Monthly means for fruit weight in Red (c and d) and Orange (e and f) cultivar are depicted in red and orange colour, respectively. Error bars indicate standard error of mean. Circles and triangles represent Autumn (AE) and Summer (SE) experiment, respectively. Control and SG treatments are depicted in solid and dashed lines, respectively. Statistical significance levels (Students *t*-test) for SG effect are shown for individual months: *, *P* ≤ 0.05; **, *P* ≤ 0.01: ***, *P* ≤ 0.001.

**Figure 5.**
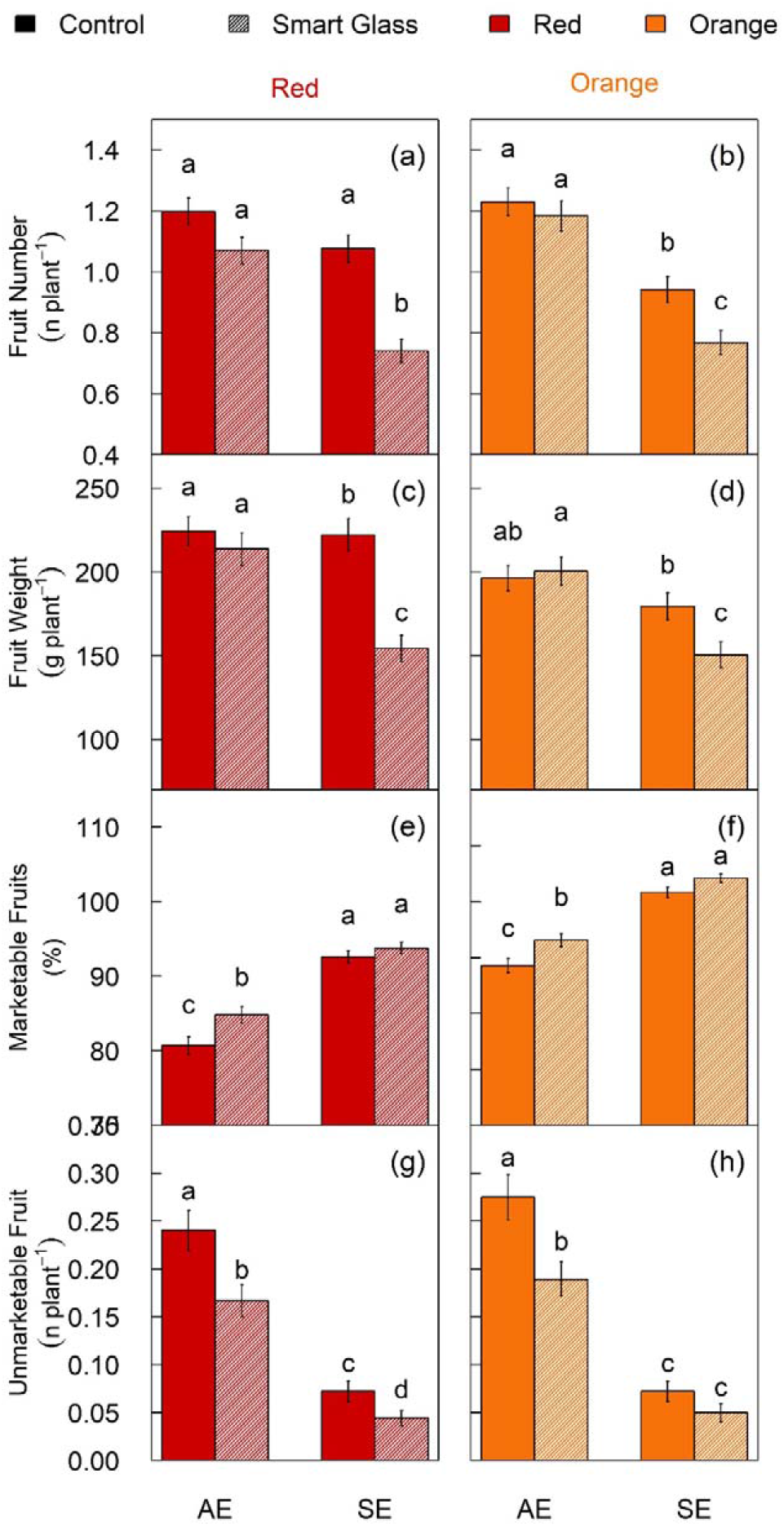
Smart Glass (SG) impact on mean harvested fruit number, fruit weight, and percentage of marketable fruits in Autumn experiment (AE) and Summer experiment (AE). Bar plots depict the mean fruit number (a and b), mean fruit weight (c and d) and percent marketable fruit (e and f) for Red and Orange cultivars, respectively. The error bars indicate the standard error (se) in each harvest month (n>120). Control and SG treatments are depicted in solid and diagonal line patterns, respectively. Bars sharing the same letter in the individual panels are not significantly different according to Tukey’s HSD test at the 5% level. Y-axis range is modified for panels c-f.

SG reduced the overall fruit number and weight, but the proportion of marketable fruits was higher in SG than in the Control. In ascending photoperiod, marketable fruit proportion increased under SG in both Red (+5.6%, *P =* 0.03) and Orange (+5.9%, *P =* 0.04) cultivars.

The proportion of marketable fruits was higher (+14%, *P* ≤ 0.001) in descending photoperiod relative to ascending photoperiod, due to a significant (50%, *P* = 0.006) reduction in small (<100 g) unmarketable fruits from both cultivars produced under SG during the descending photoperiod (Table 1). SG also reduced the number of extra-large fruits (> 250 g) during the descending photoperiod in both Red (−32 %, *P* = 0.006) and Orange cultivars (−14 %, *P* = 0.01).

In summary, SG reduced yield in SE with descending photoperiod due to light reduction under SG during initial crop growth but not in AE with ascending photoperiod involving light reduction towards the end of the growth season.

### 3.4. SG reduces net photosynthesis without altering photosynthetic capacity

SG did not affect *A*_sat_ in both cultivars across the seasonal experiments, but *A*_sat_ was significantly higher in Red cultivar relative to Orange cultivar during the descending photoperiod (Figure 6-a and b). *g_s_* was significantly higher (+42%, *P* ≤ 0.05) under SG for the Red cultivar during the ascending photoperiod (Figure 6-c). SG increased C_i_/C_a_(+7%, *P* = 0.02) and E (+31%, *P* = 0.01), but reduced PWUE (−23%, *P* ≤ 0.05) in the Red cultivar during the ascending photoperiod. PWUE in the Red cultivar, was significantly higher (*P* ≤ 0.001) due to low *g_s_* during the descending photoperiod relative to ascending photoperiod (Table 3). For light response photosynthetic parameters, SG did not significantly affect *A_max_* or *ϕ*_max_. In both experiments and cultivars, but SG reduced the θ (2 to 8%, *P* ≤ 0.05) and R_d_ (9 to 24%, *P* ≤ 0.05) (Table 2). Similar to *A*_sat_, *A*_model_ was significantly higher in Red cultivar relative to the Orange cultivar during the descending photoperiod. High *A*sat and *A*_model_ in Red cultivar indicate higher photosynthetic light use efficiency compared to the Orange cultivar. SG reduced average canopy PAR more in descending photoperiod (~−22%, *P* ≤ 0.001) than ascending photoperiod (~−18%, *P* ≤ 0.001). In ascending photoperiod, SG similarly reduced *A_model_* (~ −12%, *P* ≤ 0.001) in both cultivars, but in descending photoperiod, SG reduced *A_model_* more in Red (−17%, *P* ≤ 0.001) with higher *A_sat_* rates than in the Orange (−14%, *P* ≤ 0.001) cultivar (Figure 6-e, f, and Table 2).

**Figure 6.**
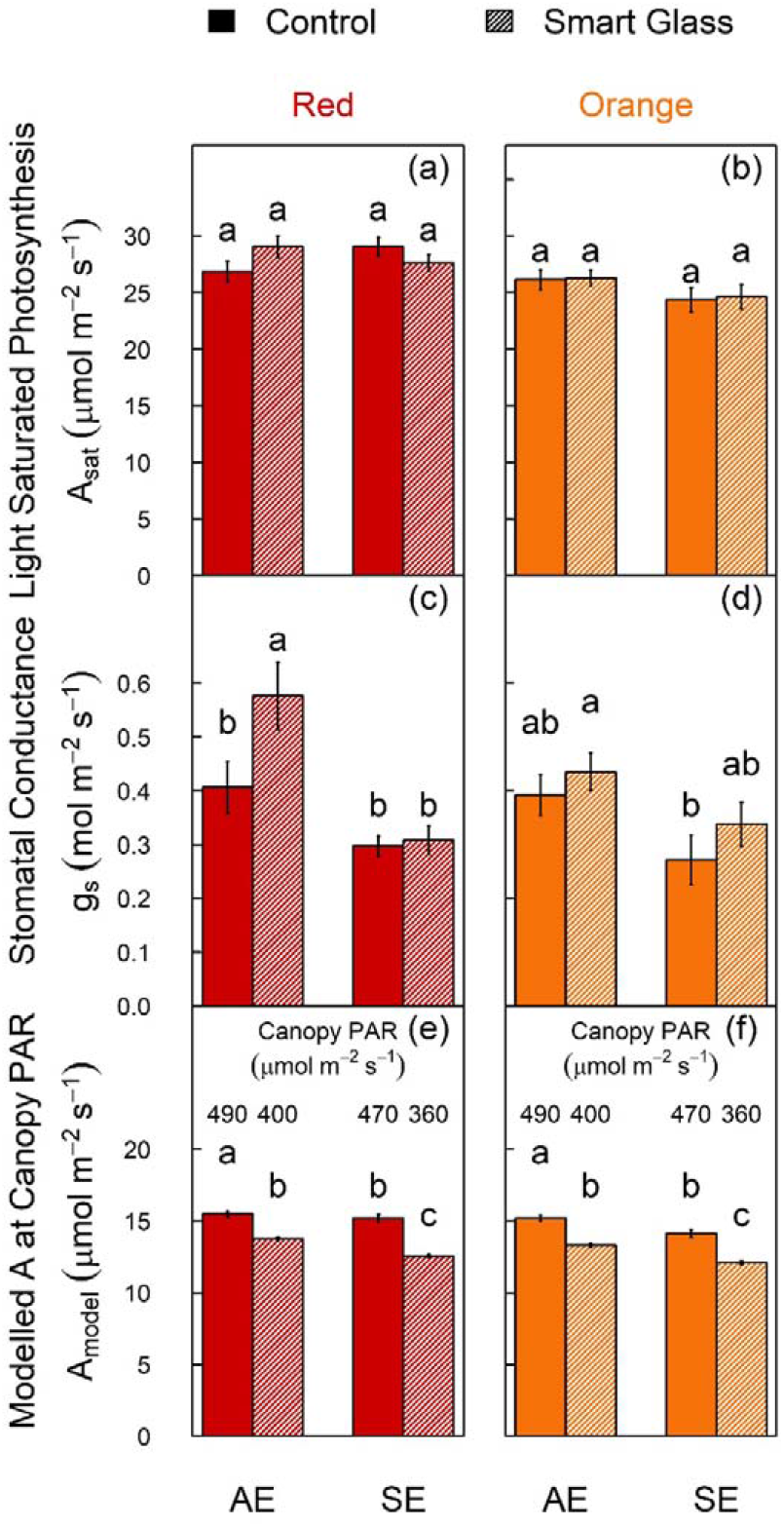
Smart Glass (SG) impact on light saturated photosynthesis (*A_sat_*), stomatal conductance (g_s_), and modelled photosynthetic rates (*A_model_*) at canopy photosynthetically active radiation (PAR). Bar plot of means for light saturated CO_2_ assimilation rate (*A_sat_*, a and b), light saturated stomatal conductance (g_s_, c and d) and modelled CO_2_ assimilation rate (*A_model_*, e and f) at the mean canopy PAR level (shown in figure) in SG and control bays, in Red and Orange capsicum cultivars. Data are shown for Autumn Experiment (AE with ascending photoperiod) and Summer Experiment (SE with descending photoperiod). Error bars indicate standard error of mean. Red and Orange cultivars are depicted in red and orange colour, respectively. Control and SG treatments are depicted in solid and diagonal line patterns, respectively. Bars sharing the same letter in the individual panels are not significantly different according to Tukey’s HSD test at the 5% level.

**Table 2.** Summary of statistical analysis for the Smart Glass (SG) and Experiment (Exp) effect on photosynthetic parameters (n=10). Data are shown as mean ± standard error. *P* values (≤ 0.05 are significant) are given according to the one-way analysis of variance (ANOVA) for SG and two-way ANOVA for SG*Exp interaction along with Exp main effects. Alternatively, Welch’s F-test for unequal variance with normal distribution or Kruskal-Wallis test for equal/unequal variance with non-normal distribution was used instead of one-way ANOVA. AE – Autumn Experiment with ascending photoperiod, SE – Sumer Experiment with descending photoperiod. NA – Not Available (as two-way ANOVA assumptions were not met).

| Parameter | Exp | Red |  | Change<br>(%) | SG<br>*P*-value | Exp<br>*P*-value | SG*Exp | Orange |  | Change<br>(%) | SG<br>*P*-value | Exp<br>*P*-value | SG*Exp |
| --- | --- | --- | --- | --- | --- | --- | --- | --- | --- | --- | --- | --- | --- |
|  |  | Control | Smart Glass |  |  |  |  | Control | Smart Glass |  |  |  |  |
| Light Saturated Photosynthesis Parameters |  |  |  |  |  |  |  |  |  |  |  |  |  |
| **A<sub>sat</sub>** (μmol m<sup>-2</sup>s<sup>-1</sup>) | AE | 26.9±0.9 | 29.1±1.0 | +8.2 | 0.11 | 0.66 | **0.04** | 26.1±0.9 | 26.3±0.7 | +0.6 | 0.89 | 0.08 | 0.95 |
|  | SE | 29.1±0.8 | 27.6±0.7 | -5.0 | 0.20 |  |  | 24.4±1.1 | 24.6±1.1 | +1.1 | 0.86 |  |  |
| **g<sub>s</sub>** (mol m<sup>-2</sup>s<sup>-1</sup>) | AE | 0.41±0.05 | 0.58±0.06 | **+41.5** | **0.02** | **NA** | NA | 0.39±0.04 | 0.43±0.04 | +10.3 | 0.41 | **0.009** | 0.76 |
|  | SE | 0.30±0.02 | 0.31±0.03 | +3.3 | 0.75 |  |  | 0.27±0.05 | 0.34±0.04 | +25.9 | 0.07 |  |  |
| **PWUE** (A<sub>sat</sub>/g<sub>s</sub>) | AE | 71.2±4.6 | 55.2±4.7 | **-22.6** | **0.02** | **6.6×10<sup>-7</sup>** | 0.3 | 70.5±4.8 | 63.9±4.9 | -9.3 | 0.35 | **0.001** | 0.27 |
|  | SE | 99.7±4.4 | 95.8±8.8 | -3.9 | 0.69 |  |  | 102.1±8.6 | 80.6±7.8 | -21.1 | 0.08 |  |  |
| **Ci/Ca** | AE | 0.71±0.02 | 0.76±0.02 | **+7.0** | **0.02** | **2.8×10<sup>-5</sup>** | 0.35 | 0.71±0.02 | 0.74±0.02 | +4.2 | 0.31 | **0.02** | 0.31 |
|  | SE | 0.64±0.02 | 0.65±0.03 | +1.6 | 0.63 |  |  | 0.63±0.03 | 0.70±0.03 | +11.1 | 0.08 |  |  |
| **Transpiration rate** (E) | AE | 4.38±0.24 | 5.77±0.41 | +31.7 | 0.01 | NA | NA | 4.93±0.19 | 4.90±0.27 | -0.6 | 0.92 | **0.03** | 0.16 |
|  | SE | 4.81±0.13 | 4.53±0.27 | -5.8 | 0.35 |  |  | 4.00±0.35 | 4.69±0.17 | +17.3 | 0.09 |  |  |
| Light Response Curve Parameters |  |  |  |  |  |  |  |  |  |  |  |  |  |
| **A<sub>max</sub>**(μmol m<sup>-2</sup>s<sup>-1</sup>) | AE | 27.6±1.5 | 31.6±1.2 | +14.4 | 0.05 | 0.5 | 0.1 | 26.8±1.2 | 27.7±0.9 | -3.3 | 0.55 | 0.01 | 0.66 |
|  | SE | 28.9±1.5 | 28.9±1.2 | +0.0 | 0.99 |  |  | 23.2±1.2 | 25.1±1.2 | +8.1 | 0.28 |  |  |
| **Theta** (θ) | AE | 0.91±0.01 | 0.84±0.01 | **-7.7** | **5.4×10<sup>-4</sup>** | 0.40 | 0.06 | 0.88±0.01 | 0.84±0.01 | **-4.5** | **0.04** | 0.24 | 0.30 |
|  | SE | 0.90±0.01 | 0.88±0.02 | -2.2 | 0.37 |  |  | 0.92±0.02 | 0.84±0.02 | **-8.7** | **0.01** |  |  |
| **R<sub>d</sub>** (μmol m<sup>-2</sup>s<sup>-1</sup>) | AE | -2.54±0.10 | -1.93±0.07 | **+24.0** | **8.7×10<sup>-5</sup>** | **0.08** | 0.14 | -2.15±0.0 | -1.78±0.10 | **+17.2** | **0.01** | **0.04** | 0.93 |
|  | SE | -2.10±0.12 | -1.90±0.20 | +9.5 | 0.40 |  |  | -1.90±0.1 | -1.55±0.09 | +18.4 | 0.05 |  |  |
| **Phi** (Φ<sub>max</sub>) | AE | 0.078±0.002 | 0.079±0.002 | 0.01 | 0.88 | **0.01** | 0.16 | 0.076±0.0 | 0.073±0.001 | -12.5 | 0.32 | 0.84 | 0.53 |
|  | SE | 0.087±0.002 | 0.081±0.001 | -11.1 | 0.07 |  |  | 0.074±0.0 | 0.074±0.002 | 0 | 0.9 |  |  |
| Photosynthesis at mean growth light - Canopy PAR between 9 am to 3 pm and modelled photosynthetic rate at canopy PAR (Amodel) |  |  |  |  |  |  |  |  |  |  |  |  |  |
| **Canopy PAR**<br>(μmol m<sup>-2</sup>s<sup>-1</sup>) | AE | 490 ± 2 | 401 ± 1 | -18 | NA | NA | NA | 490 ± 2 | 401 ± 1 | -18 | NA | NA | NA |
|  | SE | 468 ± 2 | 361 ± 1 | -22 | NA |  |  | 468 ± 2 | 361 ± 1 | -22 | NA |  |  |
| **Amodel**<br>(μmol m<sup>-2</sup>s<sup>-1</sup>) | AE | 15.5 ± 0.2 | 13.8 ± 0.1 | **-11** | **1.5×10<sup>-6</sup>** | **NA** | **NA** | 15.2 ± 0.2 | 13.3 ± 0.1 | **-12** | **1.1×10<sup>-7</sup>** | **7.6×10<sup>-7</sup>** | 0.71 |
|  | SE | 15.2 ± 0.3 | 12.6 ± 0.1 | **-17** | **1.7×10<sup>-4</sup>** |  |  | 14.1 ± 0.3 | 12.1 ± 0.1 | **-14** | **8.1×10<sup>-5</sup>** |  |  |

**Table 3.**
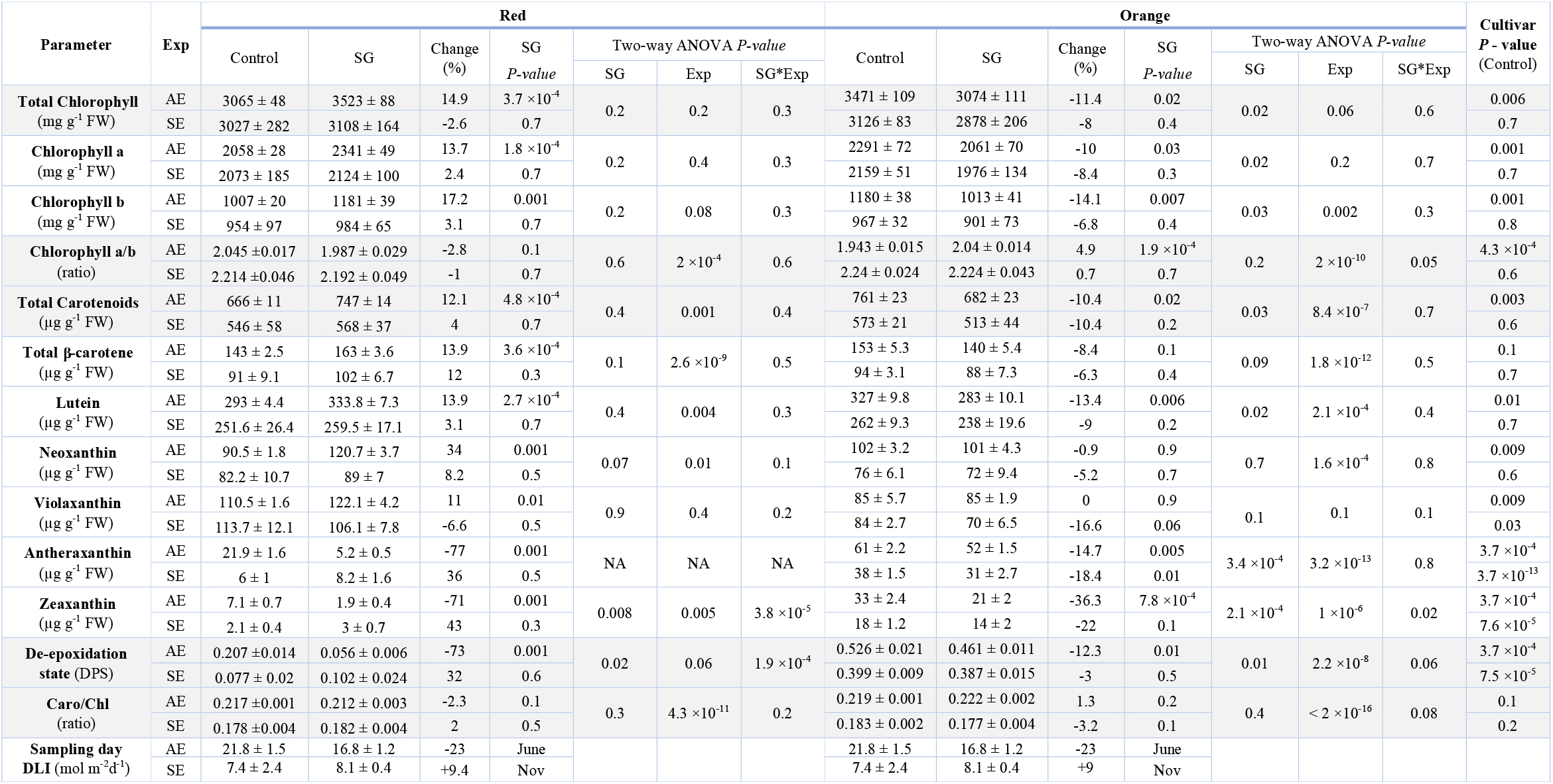
Summary of statistical analysis for the SG effect on leaf pigment composition in Autumn and summer experiment (n=9-10). The de-epoxidation state (DPS) was calculated as the ratio of (A+Z)/(A+ Z+ V), where A – antheraxanthin; Z – zeaxanthin; V – violaxanthin. Data are shown as mean ± standard error. *P* values (≤ 0.05 are significant) are given according to the one-way analysis of variance (ANOVA) for SG and two-way ANOVA for SG*Exp interaction along with Exp main effects. Alternatively, Welch’s F-test for unequal variance with normal distribution or Kruskal-Wallis test for equal/unequal variance with non-normal distribution was used instead of one-way ANOVA. Abbreviations: AE – Autumn Experiment with Ascending Photoperiod, SE – Sumer Experiment with Descending Photoperiod. NA – Not Available (as two-way ANOVA assumptions were not met).

Overall, SG did not affect photosynthetic capacity of both cultivars in AE and SE, but Red cultivar with high photosynthetic light use efficiency Relative to Orange, showed a greater reduction in net photosynthesis during descending relative to ascending photoperiod.

### 3.5. SG impacts photosynthetic pigments during ascending photoperiods

Leaf photosynthetic pigments were quantified from both cultivars at 220 DAT and 148 DAT in AE and SE, respectively. Chlorophyll a/b (Chl a/b) and carotenoid/chlorophyll (carot/chl) ratios were not affected by the SG in either cultivar during both seasons (Table 3). The chl a/b and carot/chl ratios were significantly higher and lower respectively, during the descending relative to ascending photoperiod revealing a seasonal influence.

The Red cultivar displayed a significantly higher (+15%, *P* ≤ 0.001) foliar chlorophyll content in SG-grown plants during the ascending photoperiod, but not during the descending photoperiod, relative to the Control (Figure 7 and Table 3). Similarly, SG slightly increased total carotenoid content by 12% (*P* ≤ 0.001) during the ascending photoperiod due to increased neoxanthin, violaxanthin, lutein and β-carotene. In contrast, there was a substantial reduction in antheraxanthin and zeaxanthin in SG-grown Red cultivar leaves that led to a dramatic reduction (−73%, *P* ≤ 0.001) in the DPS. At the leaf sampling stage (roughly 150 DAT), the 23% reduction in the DLI during the ascending photoperiod correlated with a lower DPS revealing that the Red cultivar has capacity to acclimate to the SG light environment during the ascending photoperiod. During the descending photoperiod the SG did not affect DLI levels at the leaf sampling stage which were already considerably lower, and individual carotenoids and the DPS remained unchanged in SG-grown plants of the Red cultivar, which were significantly lower compared to the ascending photoperiod (Figure 7 and Table 3). In summary, Red capsicum leaves from SG plants during the ascending photoperiod showed a major reduction in DPS in response to reduced DLI, and total pigment reduction. This was not found during the descending photoperiod when DLI levels were lower and unaffected by the SG (Figure 7).

**Figure 7.**
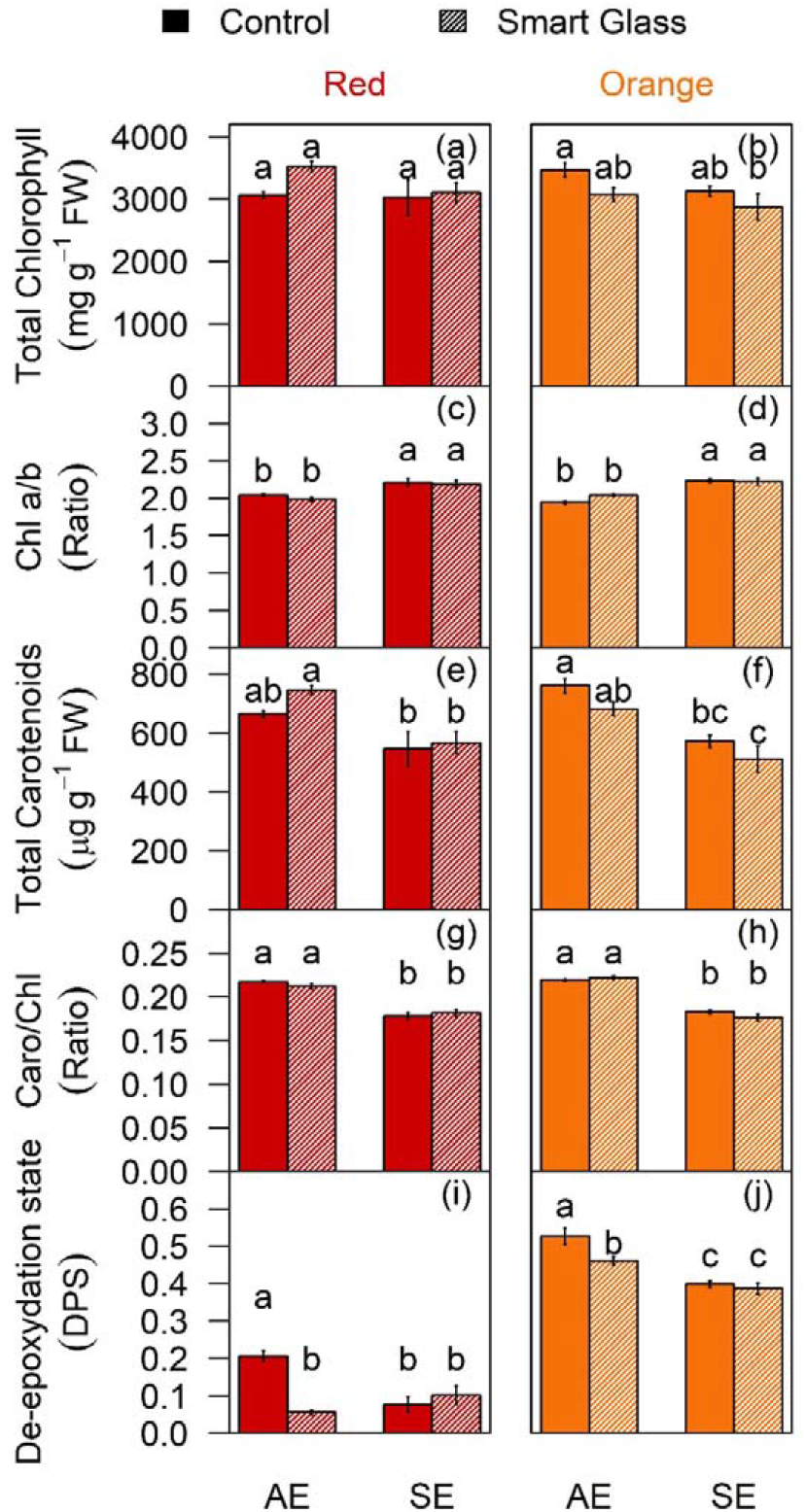
Smart Glass (SG) impact on leaf total chlorophyll and carotenoid concentration, and de-epoxidation state (DPS, unitless). Bar plots of total chlorophyll (a and b), Chlorophyll a/b (Chla/b) ratio (c and d), total carotenoids (e and f), carotenoid/chlorophyll (Caro/Chl) ratio (g and h) and DPS (i and j), are shown as means ± standard error with 8-12 biological replicates per cultivar during Autumn (AE) and summer (SE) experiments with ascending and descending photoperiod respectively. The de-epoxidation state (DPS) was calculated as the ratio of (A+Z)/(A+ Z+ V), where A – antheraxanthin; Z – zeaxanthin; V – violaxanthin. Control and SG treatments are depicted in solid and diagonal line patterns, respectively. Bars sharing the same letter in the individual panels are not significantly different according to Tukey’s HSD test at the 5% level.

In contrast to the Red cultivar, the Orange cultivar showed a significant decrease in total leaf chlorophyll (−11%, *P* = 0.02) and carotenoid (−10%, *P* = 0.02) levels when grown under the SG compared to plants in the Control during the ascending photoperiod. The SG reduced total foliar carotenoids mostly due to a 13.4, 16.6, and 18.4 % reduction in lutein, antheraxanthin, and zeaxanthin levels, respectively, compared to the Control (Table 3). As a result, the Orange SG-grown leaves displayed a 12.3% reduction in their DPS, similar to that of the Red cultivar, revealing that the SG grown plants that have lower DLI levels during the ascending photoperiod retain capacity to reduce their DPS. During the descending photoperiod, leaves from the Orange cultivar showed no change in total chlorophyll or carotenoid levels. However, violaxanthin and antheraxanthin were significantly reduced, which did not affect the DPS, although the DPS was significantly lower compared to the ascending photoperiod (Figure 7 and Table 3).

Both Red and Orange cultivar leaf tissues displayed significantly higher total carotenoids and chlorophyl levels during the ascending photoperiod that was reduced by the SG, yet during the descending photoperiod pigment levels were not altered by the SG and similar to levels quantified in leaves from the SG harvested during the ascending photoperiod. Intriguingly, the leaves of the Orange cultivar were enriched in antheraxanthin and zeaxanthin by 2-10 fold compared to the Red cultivar (Figure 7 and Table 3). Both cultivars showed that SG induces a lower DPS ratio during the ascending photoperiod that correlated with a lower DLI, and that during the descending photoperiod the overall lower DLI reduced the DPS to a threshold unaffected by the SG (Figure 7 and Table 3). Hence, the DPS and DLI may be useful indicators of seasonal effects on photosynthetic capacity of greenhouse capsicum.

### 3.6. Correlation analysis among light and plant growth traits

The positive correlation between plant height and DLI was stronger in ascending photoperiod (R^2^ = 0.7 and 0.6 in Red and Orange, respectively, *P* ≤ 0.01) relative to descending photoperiod (R^2^ = 0.4 (ns) and 0.5 (*P* ≤ 0.05) in Red and Orange respectively). The coefficient of mean describing the change in height per unit DLI was also significantly higher in ascending (>0.47) relative to descending photoperiod (0.15) in both cultivars, indicating that light use efficiency for plant growth was better during ascending photoperiod (Figure S2a-b). Moreover, the impact of SG on this relationship was stronger in descending photoperiod, which is evident from clear separation of correlation points between Control and SG. In contrast, the positive correlation between increase in plant height and photoperiod (daylength) was stronger in descending (R^2^ = 0.3 and 0.6 (*P* ≤ 0.001) in Red and Orange, respectively) relative to ascending photoperiod (R^2^ = 0.2 in only Red, *P* ≤ 0.05) (Figure S3). The number of buds and developing fruits negatively correlated only in descending photoperiod (R^2^ = 0.3, *P* ≤ 0.01), indicating fewer buds as the fruits develop, particularly during the initial growth period under high light levels and longer photoperiod (Figure S2a-b and Figure S3). Similarly, the number of buds and photoperiod negatively correlated only in SE with descending photoperiod (R^2^ = 0.7, *P* ≤ 0.001 in Red cultivar and R^2^ = 0.6,*P* ≤ 0.01 in Orange cultivar, respectively), indicating fewer buds as the day length increased (Figure S3).

Thus, increases in DLI and daylength both promoted rate of height increases but there was no correlation between height increase and daylength during ascending photoperiod as most of the early active growth occurred during relatively similar shorter photoperiods of late Autumn and Winter. Similarly, number of buds decreased with increase in developing fruits during descending photoperiod suggesting source sink regulation to maintain sustainable fruit development but there was lack of correlation between bud number and daylength, and between bud number and developing fruits in ascending photoperiod due to most of the early crop growth during relatively similar shorter photoperiods of late Autumn and Winter.

## 4. Discussion

In this study, we tested the impact of SG on two capsicum cultivars in two seasonal trials and found that the SG and time of planting affected plant traits via alterations in light quantity and quality. Firstly, SG reduced plant growth and yield traits by significantly reducing light during the descending photoperiod that progressed into winter. Secondly, time of planting affected plant growth and development as a function of DLI that correlated with the DPS as plants were exposed to low and high light levels at early growth stages during ascending and descending photoperiod experiments. Thirdly, SG reduced DLI to a greater degree during descending photoperiod (−22%) than during ascending photoperiod (−18%). Overall, SG film reduced capsicum production during the descending photoperiod, but did not affect production during the ascending photoperiod. The ability of the two capsicum cultivars to adapt to the SG-environment did not correlate with photosynthetic capacity (*A_max_/A_sat_*), nor total pigment levels or chl/carot ratios, yet the lower DPS reflected the lower DLI that associated with a reduction in yield.

### 4.1. SG reduced capsicum yield is minimized during ascending photoperiods

Greater light intensity can increase capsicum yield, such that each additional 1% of light can promote greenhouse vegetable fruit production by 0.7 - 1% under limiting-light conditions (Marcelis et al., 2006). Given that the light levels in this study were often below the minimum required DLI (15 mol m^−2^ d^−1^), the yield strongly responded to changes in light. The first harvest occurred 18 days earlier in ascending photoperiod (period of lower light and shorter photoperiod) relative to descending photoperiod (period of higher light and longer photoperiod). Both cultivars produced more fruit at the first harvest in descending photoperiod than ascending photoperiod in control and SG treatment, due to higher rates of net photosynthesis and higher translocation rates due to the longer photoperiod (Dorais et al., 1996).

Despite small differences in total DLI reduction under SG, descending showed higher and more consistent reductions in yield relative to ascending photoperiod in both cultivars, thus accepting our hypothesis that SG will reduce yield more during descending than ascending photoperiod. This indicates that high light intensity and the corresponding reduction of light under SG during early plant development had a strong negative impact on yield rather than light limitation under SG towards the end of the growth season (Figure 4, c-f). The reduction of fruit number and weight in both cultivars in descending photoperiod (Table 2) was consistent with the reduction of eggplant fruits under SG (Chavan et al., 2020). Hence, in both studies, yield reductions were likely to be driven by limited carbon supply for translocation to the fruit due to the reduced source activity (i.e., photosynthesis). A significant reduction in yield associated with decreased red and far-red light under SG is consistent with a previous studies in tomatoes (Kalaitzoglou et al., 2019) and capsicum (Tang et al., 2019) due to reduced source strength as a consequence of reduced red and far-red light, which could be explained by the additive effect of far-red light and the potential to boost photosynthesis (Zhen and Bugbee, 2020). Therefore, light reduction under SG during early crop growth in descending photoperiod reduced yield more than light reduction by SG towards the end of the growth season.

The lower or no reduction in yield under SG during ascending photoperiod, associated with Autumn planting, can be explained by initial seedling growth in shorter photoperiod with less reduction of light under SG followed by the reproductive phase with longer photoperiod with higher reduction of light under SG. This is consistent with previous studies showing best seedling growth at shorter photoperiods (Yang et al., 2017) and increases in yield due to higher translocation rates at longer photoperiods during the reproductive stage (Dorais et al., 1996). Hence, light reduction towards the end of the growing season might not have significantly impacted overall yield during ascending photoperiod.

### 4.2. DLI reduction under SG significantly decreased DPS and yield independent of impacts on photosynthetic capacity

SG grown plants exhibited reduced net photosynthesis measured at incident light evident from *A_model_*, which is consistent with our hypothesis that SG light limitation will reduce photosynthesis. However, SG did not affect the *A_sat_, A_max_* or *ϕ*_max_ (Figure 6 and Table 2) indicating no change in photosynthetic capacity. In contrast, eggplant trials under SG showed significant reduction in *A_sat_* suggesting physiological adjustment of eggplant leaf photosynthesis to an altered light environment (Chavan et al., 2020). Here, SG marginally reduced the θ in both experiments and cultivars, which indicates less agreement between light absorption and photosynthetic capacity (Evans et al., 1993). Similarly, SG reduced R_d_ in both experiments and cultivars which suggests down-regulation of respiration rates in low light (Reich et al., 1998). Therefore, SG appears to have downregulated photosynthesis due to the limited light without acclimatory changes in photosynthetic capacity associated with anatomical changes that reduce the ability to utilise additional light upon availability.

During the ascending photoperiod when DLI levels were reduced, leaves from the Orange and Red cultivars showed plasticity to change their carotenoid levels and reduce their DPS, yet during the descending photoperiod when DLI levels were lower and unchanged by the SG light environment, both cultivars did not show change in DPS as there was no need of photoprotection due to absence of high light (Figure 7). Lower light levels are generally linked to a lower DPS (Ding et al., 2006) and indeed our eggplant crop grown under SG showed a significant reduction in DPS (Chavan et al., 2020). The reduction in DPS during the eggplant trial was driven by regulation of all three xanthophyll cycle components, including Violaxanthin, Antheraxanthin and Zeaxanthin (Chavan et al., 2020). In Red and Orange capsicum cultivar leaf tissues, the changes in DPS were driven by only a reduction in Antheraxanthin and Zeaxanthin (Table 3). Unlike eggplant leaf tissues (Chavan et al., 2020), there was no reduction in carotenoid to chlorophyll ratio for capsicum cultivar leaf tissues grown under SG in the current study, yet SG induced a reduction of energy quenching antheraxanthin and zeaxanthin (DPS) in both capsicum and eggplant, indicating that xanthophyll cycle pigments provide photoprotection but they might also contribute to sustainable production of yield under favourable environmental conditions. The reduction in violaxanthin and carotenoid to chlorophyll ratio in eggplant could be linked to the photosynthetic acclimation to low light and a reduction in photosynthetic capacity in response to the SG (Chavan et al., 2020), since there was neither reduction in violaxanthin and carotenoid to chlorophyll ratio nor change in photosynthetic capacity evident from unchanged *A_sat_* or *A_max_* in the current study. In addition, a study in Arabidopsis (Kromdijk et al., 2016) has shown improved photosynthesis and crop productivity by accelerating recovery from photo-damage by increasing violaxanthin, and reducing antheraxanthin and zeaxanthin via overexpression of xanthophyll cycle enzymes.

Since the SG-induced overall reduction in DLI did not affect the pigment ratios of light harvesting complex, it is not surprising that SG foliar leaves displayed a similar photosynthetic capacity to the Control (Figure 7 and Table 3). Lower caro/chl ratio due to decrease in carotenoids, including β-carotene, neoxanthin, antheraxanthin and zeaxanthin during descending relative to ascending photoperiod, reflects less demand for light capture and photoprotection due to low light at the time of leaf sampling (Demmig-Adams et al., 1996). The precise relationship between DPS, photoperiod, and SG films is intriguing considering that both eggplant and capsicum have been similarly affected by the SG-altered light environment. The reduction in the DPS state correlated with lower yields under SG during ascending photoperiod, and the overall lower DPS during descending photoperiod was linked to lower DLI levels (at the time of sampling) indicating that carotenoid pigment ratios are tightly linked to SG-altered light levels. In summary, light limited photosynthesis and yield under SG is associated with lower DPS with or without the modification of photosynthetic capacity driven by changes in all three xanthophylls or only in antheraxanthin and zeaxanthin, respectively.

In descending photoperiod experiment, Red cultivar with higher *A_sat_* and *A_max_* relative to Orange displayed stronger reductions in *A_model_*. These reductions in *A_model_* corresponded with yield reductions in response to the SG during descending photoperiod. However, both cultivars displayed similar reduction in *A_model_*, which corresponded with no changes in yield during ascending photoperiod (Figure 6 and Table 2). Whilst most of the leaf pigments, except violaxanthin, were generally high in Orange relative to Red cultivar, the DPS was substantially higher (1-5 fold) due to the increased levels of Antheraxanthin and Zeaxanthin, suggesting a stronger capacity to dissipate excessive light as heat. However, both Antheraxanthin and Zeaxanthin were more prominently reduced in the Red cultivar that displayed more yield reductions (Figure 5). Taken together, in Control glass, Red cultivar was able to utilize the additional high light due to the high levels of light harvesting violaxanthin and reduce the energy loss via dissipation as heat due to low levels of energy quenching antheraxanthin and zeaxanthin compared to Orange cultivar. Ability to utilise excessive light could be explained by higher *A_sat_, A_max_* and consequently yield in Red compared to Orange cultivar which was more resilient to changes in yield and DPS. This is consistent with a previous Arabidopsis study showing increases in productivity by increasing violaxanthin and reducing antheraxanthin and zeaxanthin (Kromdijk et al., 2016).

### 4.3. SG affects greenhouse capsicum yield in a season and cultivar dependent manner

Consistent growth rates for fruit and vegetable production depend on the availability of natural sunlight during the growing season and the transmittance of light through glasshouse films. The general DLI recommendation for greenhouse fruit and vegetable production is > 13 mol m^−2^ d^−1^ while capsicum requires a minimum of 15 mol m^−2^ d^−1^ with a recommended DLI > 20 mol m^−2^ d^−1^ (Fan et al., 2013). In the current study, plants grown under SG were mostly light-limited except during Summer, while plants grown under Control glass were light-limited only during Winter (Figure 2). This is line with our hypothesis that SG will decrease the beneficial light in PAR region for photosynthesis and growth.

Capsicum, a non-photoperiod sensitive plant, shows best seedling growth under short photoperiods (Tang et al., 2019; Yang et al., 2017), but further vegetative and reproductive growth is better under longer photoperiods (Yamamoto et al., 2008). In the current study, changes in both photoperiod and light intensity affected plant growth traits in Red and Orange capsicum cultivars. Higher DLI with longer photoperiod (Figure 2-C) during early seedling establishment (< 10 weeks) promoted faster growth and increased plant height in descending photoperiod compared to ascending photoperiod, while lower PAR and far-red light under SG led to a marginal decrease in height in both cultivars and experiments (Figure 3a-b). Light drives photosynthesis and is a key determinant of crop growth (Tang et al., 2019), which explains the slightly reduced height in SG grown plants during the initial development due to lower light. SG also significantly reduced IR (60%) which increased the red: far-red ratio. This effect may have decreased stem elongation, and hence height, in the current study. This is consistent with a previous study in tomato (Runkle and Heins, 2002), which found that far-red deficiency reduced stem length for shade avoidance (Ballaré and Pierik, 2017; de Wit et al., 2016). Nonetheless, the decrease in height in both cultivars was marginal, and eventually SG grown plants were similar to Control plants in ascending photoperiod, but not in descending photoperiod.

High abortion of flowers and developing fruits were only observed in ascending photoperiod at 65 DAT, potentially due to a lower supply of photo-assimilates under low light, which may lead to bud, flower or fruit abortion in capsicum (Wubs et al., 2009). This also explains the significant increase in the number of buds and developing fruits per stem, while maintaining a relatively similar flower number in descending relative to ascending photoperiod. In addition, SG promoted overall seasonal mean bud number only during ascending photoperiod, which were likely the result of low light conditions during early development in both Control and SG (Table 1). Furthermore, there was a strong negative relationship between the number of buds and developing fruits, and between the number of buds and photoperiod for both cultivars in descending photoperiod (Figure S2 and S3). These relationships indicate bud abortion increased with increasing fruit set and photoperiod due to the competition between the two sinks for assimilates with more availability of light during longer photoperiods (Marcelis et al., 2004).

Cultivar specific variation of yield in response to SG in the current study is consistent with variation in yield for tomato cultivars grown under photo-selective cover materials (Loik et al., 2017), suggesting that genotype is an important factor in the regulation of flowering and fruit set, which affects fruit and seed production under cover materials (Passam and Khah, 1992). Interestingly, SG increased the proportion of marketable fruits in both experiments (Table 2), consistent with a previous study in tomatoes that reported increased marketable fruits under shading by avoiding environmental stresses such as excessive radiation and temperature (Milenković et al., 2020). Thus, use of SG coupled with Autumn or Winter planting may be beneficial in the greenhouse production of high radiation sensitive fruits.

## 5. Conclusions

SG reduced most of the heat generating long wavelengths of light which may lead to energy savings in warm periods. The reduction in PAR under SG was greater during high relative to low light conditions. Hence, greater reduction of light during early crop growth in SG resulted in lower crop yields in descending compared to ascending photoperiod. The reduction in yield under SG was driven by light-limited photosynthesis without acclimatory changes in photosynthetic capacity (*A_sat_*) and overall leaf pigments (chlorophylls, carotenoids and their ratios). However, SG did reduce DPS to a greater degree in the Red cultivar relative to Orange cultivar during an ascending photoperiod suggesting a link between reduction of yield and DPS under SG. Compared to Orange, the Red cultivar was able to use the high light more efficiently for photosynthesis and yield but at the same time was more susceptible to yield reduction associated with DPS in response to low light under SG.

We conclude that light-limited photosynthesis in SG film reduced capsicum yield, which may be partially mitigated by planting in shorter photoperiods with low light (Autumn/Winter). The impact of SG on yield varies with cultivar according to foliar pigment composition associated with light capture and non-photochemical quenching via xanthophyll cycle and DPS. The link between reduction in DPS and yield under SG is intriguing and must be explored further to be used as an indicator for cultivar monitoring and validating the impact of light altering cover films on crop yield.

## 6. Acknowledgements

We are grateful to Dr. Wei Liang and Mr Norbert Klause for crop growth and management, Dr Eric Brenya for help with HPLC pigment data collection, Professor Paul Holford, Dr Rosalie Durham and Dr. Mahnaz Shahnaseri for providing fruit quality analysis methods.

## 8. Author’s contributions

S.G.C., X.H., D.T., Z.H.C., O.G., and C.I.C. designed the research. S.G.C. and X.H. with assistance from Y.A., S.A., and C.M. collected and analysed data. S.G.C., X.H., and S.A. prepared figures. S.G.C wrote the manuscript with contribution from co-authors. S.G.C. and X.H. authors contributed equally to this work

## References

Alagoz, Y., Dhami, N., Mitchell, C., Cazzonelli, C.I., 2020. cis/trans Carotenoid Extraction, Purification, Detection, Quantification, and Profiling in Plant Tissues, in: Rodríguez-Concepción, M., Welsch, R. (Eds.), Plant and Food Carotenoids: Methods and Protocols. Springer US, New York, NY, pp. 145–163. https://doi.org/10.1007/978-1-4939-9952-1_11

Aloni, B., Karni, L., Zaidman, Z., Schaffer, A.A., 1996. Changes of Carbohydrates in Pepper (Capsicum annuum L.) Flowers in Relation to Their Abscission Under Different Shading Regimes. Ann. Bot. 78, 163–168. https://doi.org/10.1006/anbo.1996.0109

Anwar, S., Nayak, J.J., Alagoz, Y., Wojtalewicz, D., Cazzonelli, C.I., 2022. Purification and use of carotenoid standards to quantify cis-trans geometrical carotenoid isomers in plant tissues, in: Methods in Enzymology. Academic Press. https://doi.org/10.1016/bs.mie.2022.01.005

Aroca-Delgado, R., Pérez-Alonso, J., Callejón-Ferre, Á.-J., Díaz-Pérez, M., 2019. Morphology, yield and quality of greenhouse tomato cultivation with flexible photovoltaic rooftop panels (Almería-Spain). Sci. Hortic. 257, 108768. https://doi.org/10.1016/j.scienta.2019.108768

Baker, N.R., 2008. Chlorophyll Fluorescence: A Probe of Photosynthesis In Vivo. Annu. Rev. Plant Biol. 59, 89–113. https://doi.org/10.1146/annurev.arplant.59.032607.092759

Ballaré, C.L., Pierik, R., 2017. The shade-avoidance syndrome: multiple signals and ecological consequences. Plant Cell Environ. 40, 2530–2543. https://doi.org/10.1111/pce.12914

Cerny, T.A., Faust, J.E., Layne, D.R., Rajapakse, N.C., 2003. Influence of Photoselective Films and Growing Season on Stem Growth and Flowering of Six Plant Species. J. Am. Soc. Hortic. Sci. 128, 486–491. https://doi.org/10.21273/JASHS.128.4.0486

Chavan, S.G., Maier, C., Alagoz, Y., Filipe, J.C., Warren, C.R., Lin, H., Jia, B., Loik, M.E., Cazzonelli, C.I., Chen, Z.H., Ghannoum, O., Tissue, D.T., 2020. Light-limited photosynthesis under energy-saving film decreases eggplant yield. Food Energy Secur. Early View, e245. https://doi.org/10.1002/fes3.245

Davis, P.A., Burns, C., 2016. Photobiology in protected horticulture. Food Energy Secur. 5, 223–238. https://doi.org/10.1002/fes3.97

de Wit, M., Galvão, V.C., Fankhauser, C., 2016. Light-Mediated Hormonal Regulation of Plant Growth and Development. Annu. Rev. Plant Biol. 67, 513–537. https://doi.org/10.1146/annurev-arplant-043015-112252

Demmig-Adams, B., Garab, G. III, W.W.A., Govindjee (Eds.), 2014. Non-Photochemical Quenching and Energy Dissipation in Plants, Algae and Cyanobacteria, Advances in Photosynthesis and Respiration. Springer Netherlands. https://doi.org/10.1007/978-94-017-9032-1

Demmig-Adams, B., Gilmore, A.M., Adams, W.W., 1996. Carotenoids 3: in vivo function of carotenoids in higher plants. FASEB J. 10, 403–412. https://doi.org/10.1096/fasebj.10.4.8647339

Dhami, N., Cazzonelli, C.I., 2020. Environmental impacts on carotenoid metabolism in leaves. Plant Growth Regul. 455–477. https://doi.org/10.1007/s10725-020-00661-w

Dhami, N., Drake, J.E., Tjoelker, M.G., Tissue, D.T., Cazzonelli, C.I., 2020. An extreme heatwave enhanced the xanthophyll de-epoxidation state in leaves of Eucalyptus trees grown in the field. Physiol. Mol. Biol. Plants. https://doi.org/10.1007/s12298-019-00729-6

Ding, L., Wang, K.J., Jiang, G.M., Li, Y.G., Jiang, C.D., Liu, M.Z., Niu, S.L., Peng, Y., 2006. Diurnal variation of gas exchange, chlorophyll fluorescence, and xanthophyll cycle components of maize hybrids released in different years. Photosynthetica 44, 26–31. https://doi.org/10.1007/s11099-005-0154-3

Dorais, M., Yelle, S., Gosselin, A., 1996. Influence of extended photoperiod on photosynthate partitioning and export in tomato and pepper plants. N. Z. J. Crop Hortic. Sci. 24, 29–37. https://doi.org/10.1080/01140671.1996.9513932

Dou, H., Niu, G., Gu, M., Masabni, J.G., 2017. Effects of Light Quality on Growth and Phytonutrient Accumulation of Herbs under Controlled Environments. Horticulturae 3, 36. https://doi.org/10.3390/horticulturae3020036

Elkins, C., Iersel, M.W. van, 2020. Longer Photoperiods with the Same Daily Light Integral Improve Growth of Rudbeckia Seedlings in a Greenhouse. HortScience 55, 1676–1682. https://doi.org/10.21273/HORTSCI15200-20

Evans, J., Jakobsen, I., Ögren, E., 1993. Photosynthetic light-response curves. Planta 189, 191–200. https://doi.org/10.1007/BF00195076

Fan, X.-X., Xu, Z.-G., Liu, X.-Y., Tang, C.-M., Wang, L.-W., Han, X., 2013. Effects of light intensity on the growth and leaf development of young tomato plants grown under a combination of red and blue light. Sci. Hortic. 153, 50–55. https://doi.org/10.1016/j.scienta.2013.01.017

Garcia-Plazaola, J.I., Faria, T., Abadia, J., Chaves, M.M., Pereira, J.S., 1997. Seasonal changes in xanthophyll composition and photosynthesis of cork oak (Quercus suber L.) leaves under mediterranean climate. J. Exp. Bot. 48, 1667–1674. https://doi.org/10.1093/jxb/48.9.1667

Hao, X., Papadopoulos, A.P., 1999. Effects of supplemental lighting and cover materials on growth, photosynthesis, biomass partitioning, early yield and quality of greenhouse cucumber. Sci. Hortic. 80, 1–18. https://doi.org/10.1016/S0304-4238(98)00217-9

He, X., Chavan, S.G., Hamoui, Z., Maier, C., Ghannoum, O., Chen, Z.-H., Tissue, D.T., Cazzonelli, C.I., 2022. Smart Glass Film Reduced Ascorbic Acid in Red and Orange Capsicum Fruit Cultivars without Impacting Shelf Life. Plants 11, 985. https://doi.org/10.3390/plants11070985

He, X., Maier, C., Chavan, S.G., Zhao, C.-C., Alagoz, Y., Cazzonelli, C., Ghannoum, O., Tissue, D.T., Chen, Z.-H., 2021. Light-altering cover materials and sustainable greenhouse production of vegetables: a review. Plant Growth Regul. https://doi.org/10.1007/s10725-021-00723-7

Jones, M.A., 2018. Using light to improve commercial value. Hortic. Res. 5, 1–13. https://doi.org/10.1038/s41438-018-0049-7

Kalaitzoglou, P., van Ieperen, W., Harbinson, J., van der Meer, M., Martinakos, S., Weerheim, K., Nicole, C.C.S., Marcelis, L.F.M., 2019. Effects of Continuous or End-of-Day Far-Red Light on Tomato Plant Growth, Morphology, Light Absorption, and Fruit Production. Front. Plant Sci. 10, 322. https://doi.org/10.3389/fpls.2019.00322

Kotilainen, T., Aphalo, PJ., Brelsford, CC., Böök, H., Devraj, S., Heikkilä, A., Hernández, R., Kylling, A., Lindfors, AV., Robson, TM., 2020. Patterns in the spectral composition of sunlight and biologically meaningful spectral photon ratios as affected by atmospheric factors. Agric. For. Meteorol. 291, 108041. https://doi.org/10.1016/j.agrformet.2020.108041

Kromdijk, J., Głowacka, K., Leonelli, L., Gabilly, S.T., Iwai, M., Niyogi, K.K., Long, S.P., 2016. Improving photosynthesis and crop productivity by accelerating recovery from photoprotection. Science 354, 857–861. https://doi.org/10.1126/science.aai8878

Lemarié, S., Guérin, V., Sakr, S., Jouault, A., Caradeuc, M., Cordier, S., Guignard, G., Gardet, R., Bertheloot, J., Demotes-Mainard, S., Proost, K., Peilleron, F., 2019. Impact of innovative optically active greenhouse films on melon, watermelon, raspberry and potato crops. Acta Hortic. 191–200. https://doi.org/10.17660/ActaHortic.2019.1252.25

Li, A., Li, S., Wu, X., Lu, H., Huang, M., Gu, R., Wei, L., He, A., 2015. Influence of Light Intensity on the Yield and Quality of *Houttuynia cordata*. Plant Prod. Sci. 18, 522–528. https://doi.org/10.1626/pps.18.522

Lin, T., Goldsworthy, M., Chavan, S., Liang, W., Maier, C., Ghannoum, O., Cazzonelli, C.I., Tissue, D.T., Lan, Y.-C., Sethuvenkatraman, S., Lin, H., Jia, B., Chen, Z.-H., 2022. A novel cover material improves cooling energy and fertigation efficiency for glasshouse eggplant production. Energy 251, 123871. https://doi.org/10.1016/j.energy.2022.123871

Loik, M.E., Carter, S.A., Alers, G., Wade, C.E., Shugar, D., Corrado, C., Jokerst, D., Kitayama, C., 2017. Wavelength-Selective Solar Photovoltaic Systems: Powering Greenhouses for Plant Growth at the Food-Energy-Water Nexus. Earths Future 5, 1044–1053. https://doi.org/10.1002/2016EF000531

Ma, Y.T., Wubs, A.M., Mathieu, A., Heuvelink, E., Zhu, J.Y., Hu, B.G., Cournède, P.H., de Reffye, P., 2011. Simulation of fruit-set and trophic competition and optimization of yield advantages in six Capsicum cultivars using functional–structural plant modelling. Ann. Bot. 107, 793–803. https://doi.org/10.1093/aob/mcq223

Marcelis, L.F.M., Broekhuijsen, A.G.M., Meinen, E., Nijs, E.M.F.M., Raaphorst, M.G.M., 2006. QUANTIFICATION OF THE GROWTH RESPONSE TO LIGHT QUANTITY OF GREENHOUSE GROWN CROPS. Acta Hortic. 97–104. https://doi.org/10.17660/ActaHortic.2006.711.9

Marcelis, L.F.M., Heuvelink, E., Baan Hofman-Eijer, L.R., Den Bakker, J., Xue, L.B., 2004. Flower and fruit abortion in sweet pepper in relation to source and sink strength. J. Exp. Bot. 55, 2261–2268. https://doi.org/10.1093/jxb/erh245

Milenković, L., Mastilović, J., Kevrešan, Ž., Bajić, A., Gledić, A., Stanojević, L., Cvetković, D., Šunić, L., Ilić, Z.S., 2020. Effect of shading and grafting on yield and quality of tomato. J. Sci. Food Agric. 100, 623–633. https://doi.org/10.1002/jsfa.10057

Ntinas, G.K., Link to external site, this link will open in a new window, Tsivelika, N., Krommydas, K., Kalivas, A., Ralli, P., Link to external site, this link will open in a new window, Irakli, M., 2019. Performance and Hydroponic Tomato Crop Quality Characteristics in a Novel Greenhouse Using Dye-Sensitized Solar Cell Technology for Covering Material. Horticulturae 5, 42. http://dx.doi.org.ezproxy.uws.edu.au/10.3390/horticulturae5020042

Ögren, E., Evans, J.R., 1993. Photosynthetic light-response curves. Planta 189, 182–190. https://doi.org/10.1007/BF00195075

O’Sullivan, C.A., Bonnett, G.D., McIntyre, C.L., Hochman, Z., Wasson, A.P., 2019. Strategies to improve the productivity, product diversity and profitability of urban agriculture. Agric. Syst. 174, 133–144. https://doi.org/10.1016/j.agsy.2019.05.007

Passam, H.C., Khah, E.M., 1992. Flowering, fruit set and fruit and seed development in two cultivars of aubergine (Solanum melongena L.) grown under plastic cover. Sci. Hortic. 51, 179–185. https://doi.org/10.1016/0304-4238(92)90117-U

Reich, P.B., Walters, M.B., Tjoelker, M.G., Vanderklein, D., Buschena, C., 1998. Photosynthesis and respiration rates depend on leaf and root morphology and nitrogen concentration in nine boreal tree species differing in relative growth rate. Funct. Ecol. 12, 395–405. https://doi.org/10.1046/j.1365-2435.1998.00209.x

Runkle, E.S., Heins, R.D., 2002. Stem extension and subsequent flowering of seedlings grown under a film creating a far-red deficient environment. Sci. Hortic. 96, 257–265. https://doi.org/10.1016/S0304-4238(02)00055-9

Shen, L., Lou, R., Park, Y., Guo, Y., Stallknecht, E.J., Xiao, Y., Rieder, D., Yang, R., Runkle, E.S., Yin, X., 2021. Increasing greenhouse production by spectral-shifting and unidirectional light-extracting photonics. Nat. Food 2, 434–441. https://doi.org/10.1038/s43016-021-00307-8

Shen, Y., Wei, R., Xu, L., 2018. Energy Consumption Prediction of a Greenhouse and Optimization of Daily Average Temperature. Energies 11, 65. https://doi.org/10.3390/en11010065

Smith, H.L., McAusland, L., Murchie, E.H., 2017. Don’t ignore the green light: exploring diverse roles in plant processes. J. Exp. Bot. 68, 2099–2110. https://doi.org/10.1093/jxb/erx098

Tang, Z., Yu, J., Xie, J., Lyu, J., Feng, Z., Dawuda, M.M., Liao, W., Wu, Y., Hu, L., 2019. Physiological and Growth Response of Pepper (Capsicum annum L.) Seedlings to Supplementary Red/Blue Light Revealed through Transcriptomic Analysis. Agronomy 9, 139. https://doi.org/10.3390/agronomy9030139

Timmermans, G.H., Hemming, S., Baeza, E., Thoor, E.A.J. van, Schenning, A.P.H.J., Debije, M.G., 2020. Advanced Optical Materials for Sunlight Control in Greenhouses. Adv. Opt. Mater. 8, 2000738. https://doi.org/10.1002/adom.202000738

Tollefson, J., 2022. What the war in Ukraine means for energy, climate and food. Nature 604, 232–233. https://doi.org/10.1038/d41586-022-00969-9

Turner, A.D., Wien, H.C., 1994. Photosynthesis, Dark Respiration and Bud Sugar Concentrations in Pepper Cultivars Differing in Susceptibility to Stress-induced Bud Abscission. Ann. Bot. 73, 623–628. https://doi.org/10.1006/anbo.1994.1078

Wubs, A.M., Heuvelink, E., Marcelis, L.F.M., 2009. Abortion of reproductive organs in sweet pepper (*Capsicum annuum* L.): a review. J. Hortic. Sci. Biotechnol. 84, 467–475. https://doi.org/10.1080/14620316.2009.11512550

Xu, J., Lv, Y., Liu, X., Wei, Q., Qi, Z., Yang, S., Liao, L., 2019. A general non-rectangular hyperbola equation for photosynthetic light response curve of rice at various leaf ages. Sci. Rep. 9, 1–8. https://doi.org/10.1038/s41598-019-46248-y

Yamamoto, S., Misumi, M., Nawata, E., 2008. Effects of Photoperiod on Vegetative Growth, Flowering and Fruiting of Capsicum frutescens L. and C. annuumL. in Japan. Environ. Control Biol. 46, 39–47. https://doi.org/10.2525/ecb.46.39

Yang, Z., He, W., Mou, S., Wang, X., Chen, D., Hu, X., Chen, L., Bai, J., 2017. Plant growth and development of pepper seedlings under different photoperiods and photon flux ratios of red and blue LEDs 8. https://doi.org/10.11975/j.issn.1002-6819.2017.17.023

Zhao, C., Chavan, S., He, X., Zhou, M., Cazzonelli, C.I., Chen, Z.-H., Tissue, D.T., Ghannoum, O., 2021. Smart glass impacts stomatal sensitivity of greenhouse Capsicum through altered light. J. Exp. Bot. 72, 3235–3248. https://doi.org/10.1093/jxb/erab028

Zhen, S., Bugbee, B., 2020. Far□red photons have equivalent efficiency to traditional photosynthetic photons: Implications for redefining photosynthetically active radiation. Plant Cell Environ. pce.13730. https://doi.org/10.1111/pce.13730

