## Supplementary Figures for "An energy-saving glasshouse film reduces seasonal, and cultivar dependent Capsicum yield due to light limited photosynthesis"

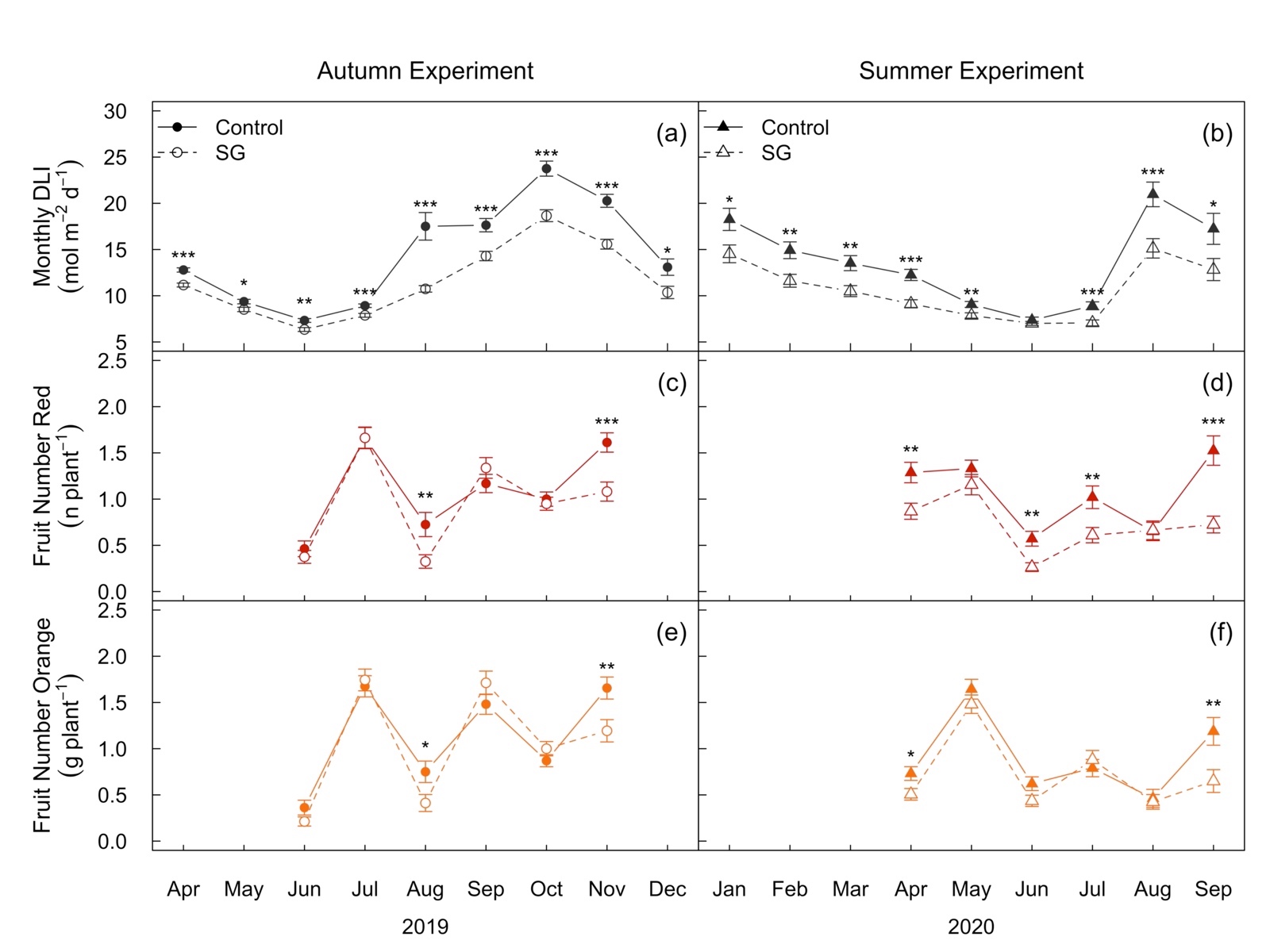

**Figure S1. Impact of Smart Glass (SG) on monthly averages of daily light integrals (DLI)and harvested fruit number across the growth season.** Panel a and b depict the monthly means for canopy level DLI during Autumn Experiment (AE) with ascending photoperiod and Summer Experiment (SE) with descending photoperiod, respectively. Monthly means for fruit number in Red (c and d) and Orange (e and f) cultivar are depicted in red and orange colour, respectively. Error bars indicate standard error of mean. Control and SG treatments are depicted in solid and dashed lines, respectively. Statistical significance levels (Students *t*-test) for SG effect are shown for individual months: *, *P* ≤ 0.05; **, *P* ≤ 0.01: ***, *P* ≤ 0.001.

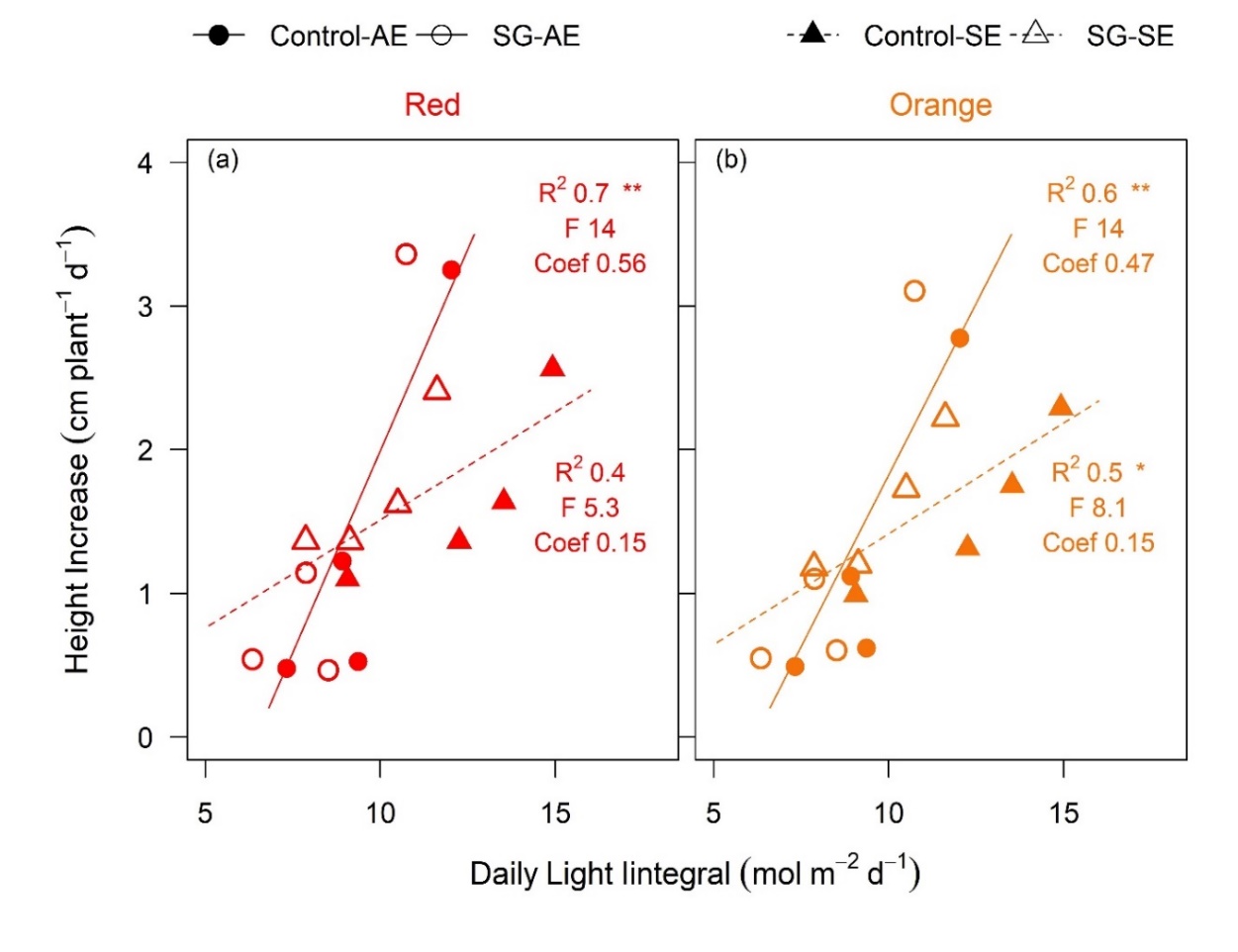

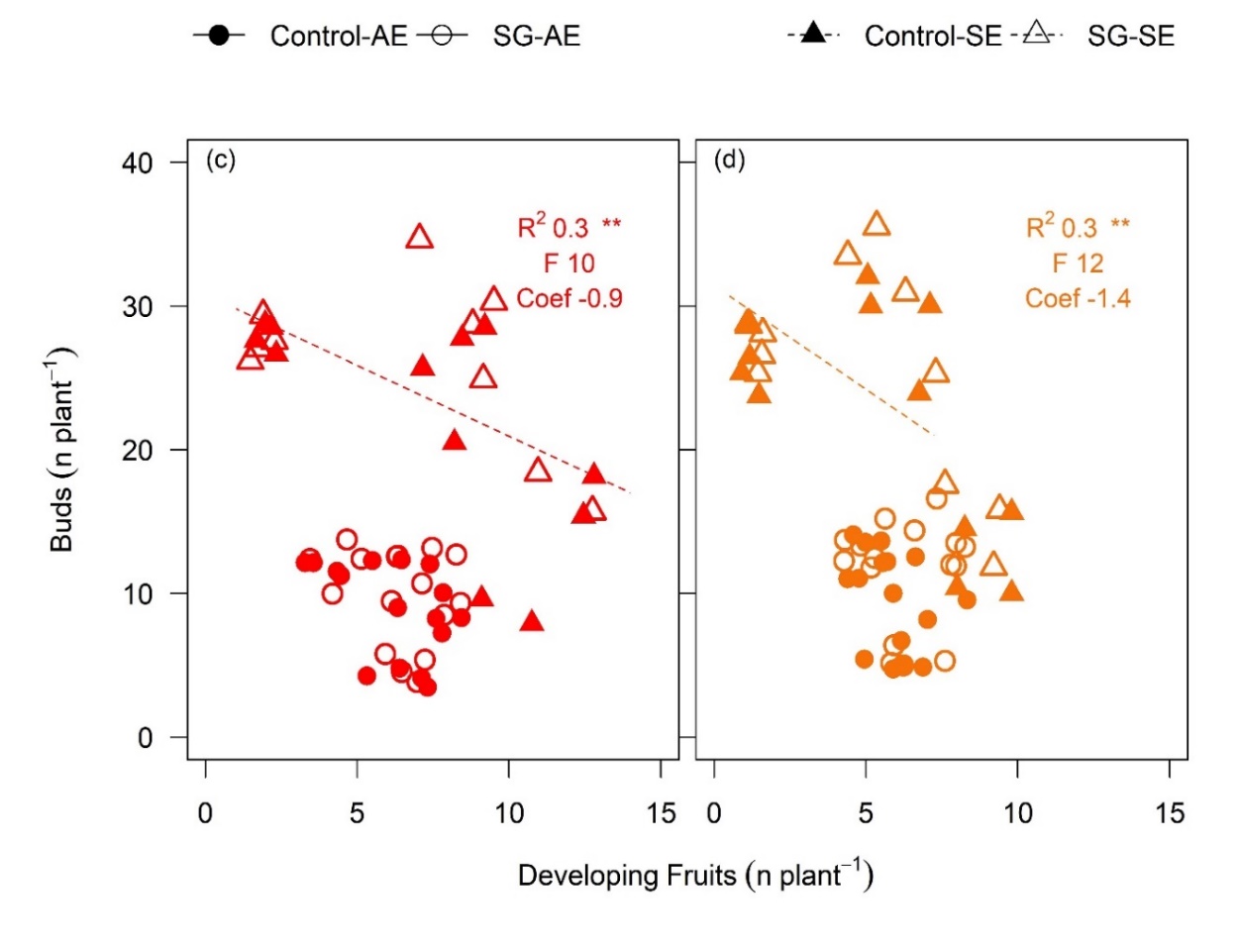

**Figure S2. SG impact on correlations between daily light integrals (DLI)and height, and bud number and developing fruit number.** Correlation between monthly means for daily height increase and canopy level DLI (a and b), and buds and developing fruits (c and d) during two experiments in Red and Orange cultivar, respectively. Red and Orange cultivars are depicted in red and orange colour, respectively. Circles with solid lines represent Autumn Experiment (AE with ascending photoperiod) and triangles with dashed line represent Summer Experiment (SE with descending photoperiod). Control and SG treatments are depicted in solid and open symbols, respectively. R^2^, F statistic and coefficients for each correlation are shown in each figure. Statistical significance levels are: * *P* ≤ 0.05; ** *P* ≤ 0.01: *** *P* ≤ 0.001.

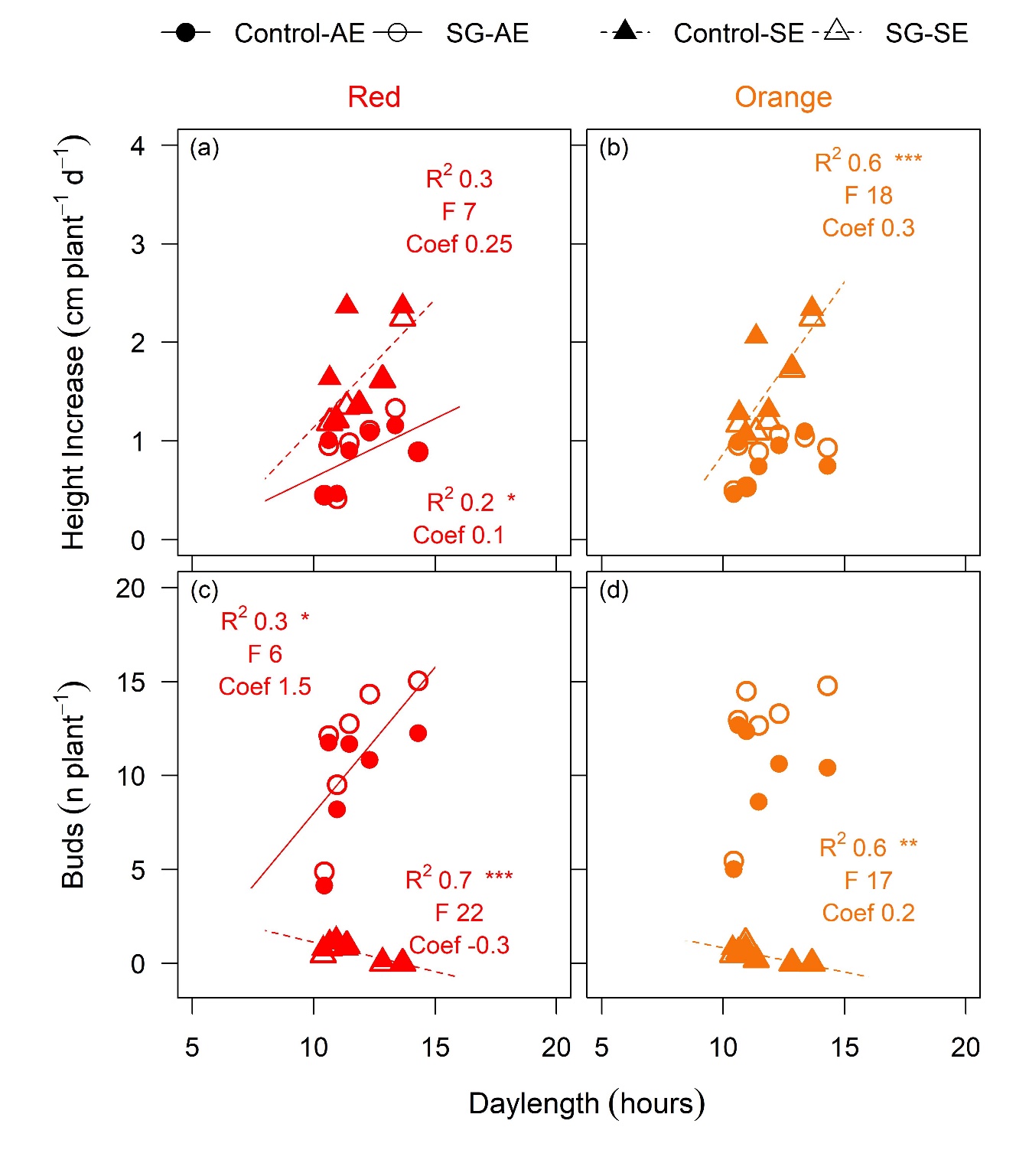

**Figure S3. SG impact on correlation of daylength with height and bud number in Autumn (AE) and Summer (SE) experiment.** Correlation between monthly means for daily height increase and day length (a and b), and bud number and daylength (c and d) during two experiments in Red and Orange cultivars, respectively. Red and Orange cultivars are depicted in red and orange colour, respectively. Circles with solid lines represent AE and triangles with dashed line represent SE. Control and SG treatments are depicted in solid and open symbols, respectively. R^2^ and *P* values for each correlation is shown.
