## Supplementary Figures for "An energy-saving glasshouse film reduces seasonal, and cultivar dependent Capsicum yield due to light limited photosynthesis"

**Table S1. Effect of Smart Glass** (**SG) on light spectrum and quality measured using radio-spectrophotometer in Autumn Experiment (AE) with ascending photoperiod.** Data are shown as mean ± standard error. *P* values are given according to the one-way analysis of variance (ANOVA) or Welch’s F-test for equal and unequal variance respectively and *P* ≤ 0.05 are significant. For non- normal distribution Kruskal-Wallis non-parametric test was used.

| Light Spectrum  (μmol m^−2^ s^−1^) | Treatment | | Change  (%) | one-way ANOVA |
| --- | --- | --- | --- | --- |
|  | Control | SG |  |  |
| UV  (221–400 nm) | 35 ± 4 | 11 ± 1 | -69 | 5.4×10^-6^ |
| Blue  (400–499 nm) | 199 ± 14 | 172 ± 4 | -14 | 0.07 |
| Green  (500–599 nm) | 279 ± 20 | 236 ± 6 | -15 | 0.05 |
| Red  (600–700 nm) | 301 ± 28 | 221 ± 7 | -26 | 0.01 |
| PAR  (400–700 nm) | 782 ± 62 | 632 ± 18 | -19 | 0.19 |
| Far-red  (700–800 nm) | 235 ± 29 | 114 ± 7 | -51 | 0.05 |

**Table S2. Effect of SG on daily light integral (DLI) during high (Autumn, Spring and Summer) and low (Winter) light periods of Autumn Experiment (AE) with ascending photoperiod and Summer Experiment (SE) with descending photoperiod.** Data are shown as mean ± standard error. *P* values are given according to the one-way analysis of variance (ANOVA) or Welch’s F-test for equal and unequal variance respectively and *P* ≤ 0.05 are significant. For non- normal distribution Kruskal-Wallis non-parametric test was used.

| Light Period | Seasons | Experiment | Mean Daily Light Integral  (DLI, mol m^-2^d^-1^) | | Change (%) | one-way ANOVA  *P* value |
| --- | --- | --- | --- | --- | --- | --- |
|  |  |  | **Control** | **SG** |  |  |
| High | Autumn, Spring and Summer | AE (2019) | 19.1 ± 0.4 | 14.8 ± 0.3 | -22 | 1.2 × 10^-13^ |
|  |  | SE (2020) | 15.9 ± 0.5 | 11.7 ± 0.3 | -26 | 3 × 10^-10^ |
| Low | Winter | AE (2019) | 8.5 ± 0.1 | 8.1 ± 0.1 | -4 | 0.02 |
|  |  | SE (2020) | 8.1 ± 0.2 | 7.4 ± 0.1 | -8 | 0.004 |
